# The Effects of Genotype × Phenotype Interactions on Silver Nanoparticle Toxicity in Organotypic Cultures of Murine Tracheal Epithelial Cells

**DOI:** 10.1101/722009

**Authors:** Tyler P. Nicholas, Anoria K. Haick, Tomomi W. Workman, William C. Griffith, James D. Nolin, Terrance J. Kavanagh, Elaine M. Faustman, William A. Altemeier

## Abstract

Silver nanoparticles (AgNP) are used in multiple applications but primarily in the manufacturing of antimicrobial products. Previous studies have identified AgNP toxicity in airway epithelial cells, but no *in vitro* studies to date have used organotypic cultures as a high-content *in vitro* model of the conducting airway to characterize the effects of interactions between host genetic and acquired factors, or gene × phenotype interactions (G×P), on AgNP toxicity. In the present study, we derived organotypic cultures from primary murine tracheal epithelial cells (MTEC) to characterize nominal and dosimetric dose-response relationships for AgNP-induced barrier dysfunction, glutathione (GSH) depletion, reactive oxygen species (ROS) production, lipid peroxidation, and cytotoxicity across two genotypes (A/J and C57BL/6J mice), two phenotypes (“Normal” and “Type 2 [T2]-Skewed”), and two exposures (an acute exposure of 24 h and a subacute exposure of 4 hours, every other day, over 5 days [5×4 h]). We characterized the “T2-Skewed” phenotype as an *in vitro* model of chronic respiratory diseases, which was marked by increased sensitivity to AgNP-induced barrier dysfunction, GSH depletion, ROS production, lipid peroxidation, and cytotoxicity, suggesting that asthmatics are a sensitive population to AgNP exposures in occupational settings. This also suggests that exposure limits, which should be based upon the most sensitive population, should be derived using *in vitro* and *in vivo* models of chronic respiratory diseases. This study highlights the importance of considering dosimetry as well as G×P effects when screening and prioritizing potential respiratory toxicants. Such *in vitro* studies can be used to inform regulatory policy aimed at special protections for all populations.

## Introduction

There is presently a need for *in vitro* models to provide relevant data on risk assessment for occupational exposures. In recent years, there has been a gradual paradigm shift toward the use of *in vitro* models that reduce, replace, and refine the use of animals in toxicity testing (the three R’s) from both the standpoint of economic imperative and animal welfare.^1,2^ Despite these economic advantages over *in vivo* models, *in vitro* models are currently limited by: 1) the use of immortalized or transformed cell lines that may not recapitulate primary cell phenotypes, 2) the use of cells from a single genetic background or phenotype, which may not capture the impact of underlying genetics or diseases on exposure risk and response variability, and 3) the limited efforts to relate *in vitro* exposures to occupational exposures in order to provide a regulatory basis for sensitivity to occupational exposures and inform regulatory policy aimed at special protections for all populations.^3,4^

Chronic respiratory diseases, including asthma, acute bronchitis, and chronic obstructive pulmonary disease (COPD), collectively affect 16% of the United States population; these diseases impair host defense mechanisms, such as barrier function and immune regulation,^5^ mucocilliary clearance and permeability,^6–12^ as well as enzymatic and non-enzymatic regulation of oxidative stress,^13,14^ and may increase an individual’s sensitivity to occupational exposures.^15–17^ Chronic respiratory diseases place a major burden on individuals, their workplaces, and the healthcare system, and yet less than 1% of the chemicals regulated by the U.S. Environmental Protection Agency’s (US EPA) Toxic Substances Control Act (TSCA) have been tested for respiratory toxicity.^3,18^ Together, the above limitations highlight a need to develop *in vitro* models suitable for high-throughput screening that also capture determinants of sensitivity in order to screen and prioritize potential respiratory toxicants while providing relevant data on risk assessment for occupational exposures.^19,20^

Silver nanoparticles (AgNP) have been tested for respiratory toxicity in previous studies, as they are used in the manufacturing of antimicrobial consumer products, including air filters, humidifiers, and purifiers as well as antimicrobial sprays.^21–23^ Previous *in vitro* studies observed the mode of action (MoA) for their antimicrobial properties (e.g., release of bioactive silver ions [Ag^+^] upon dissolution^24–26^) is also a defining factor of AgNP toxicity in mammalian cells.^27^ The MoA in airway epithelial cells is Ag^+^-mediated reactive oxygen species (ROS) production, which leads to adverse cellular responses including oxidative stress, mitochondrial dysfunction, inflammation, and cytotoxicity.^28–37^ Asthmatics are possibly a sensitive population to AgNP exposures in occupational settings, as these adverse cellular responses may further modulate innate and adaptive immune responses at the epithelial-immune interface to exacerbate this chronic airway disease.^38^

Few previous studies have characterized the effects of interactions between host genetic and acquired factors, or gene × environment interactions (G×E),^39,40^ on AgNP toxicity. Understanding G×E effects is important for identifying sensitive populations, whose underlying genetics or diseases directly modify their exposure risk and response variability to AgNP toxicity. Typically, these studies have used young, healthy animals or cell lines cultured toward a “Normal” phenotype and thus did not address the possibility of increased AgNP toxicity in asthmatics. The present study was designed to test the hypothesis that genotype and phenotype (physiological environment) will define G×E, or genotype × phenotype interaction (G×P), effects on AgNP toxicity.

We derived organotypic cultures from primary murine tracheal epithelial cells (MTEC) of differentially sensitive genotypes (A/J and C57BL/6J mice) to model the effects of host genetic factors on AgNP toxicity.^41–43^ We then differentiated these organotypic cultures toward either “Normal” or “Type 2 [T2]-Skewed” phenotypes to model the effects of host acquired factors on AgNP toxicity. To acquire the “Normal” phenotype, we used a defined differentiation media to achieve differentiation toward the cell populations in the proximal region of the conducting airway, including basal, ciliated, club, and mucin cells.^44^ To acquire the “T2-Skewed” phenotype, we used a defined differentiation media supplemented with IL-13 to skew differentiation toward an *in vitro* model of chronic respiratory diseases, characterized by barrier dysfunction,^45^ mucus production,^46^ allergic responses,^46^ and T2 responses.^47–49^

Using this high-content *in vitro* model of the conducting airway, we characterized nominal and dosimetric dose-response relationships for AgNP-induced barrier dysfunction, GSH depletion, ROS production, lipid peroxidation, and cytotoxicity across genotypes, phenotypes, and exposures to understand G×P effects on AgNP toxicity. To our knowledge, this is the first study to use organotypic cultures as a high-content *in vitro* model of the conducting airway to characterize G×P effects on AgNP toxicity. By pairing organotypic cultures with dosimetry, we can define the basis for physiologically-relevant dose-response relationships by accounting for association and dissolution, and by using a benchmark dose (BMD) approach, we can identify the most sensitive adverse cellular responses to help define a regulatory basis for G×P effects on AgNP toxicity.

## Materials and Methods

### Cell culture

All animal studies were approved by the Institutional Animal Care and Use Committee at the University of Washington. We harvested tracheas from A/J and C57BL/6J mice (The Jackson Laboratory, Bar Harbor, ME, USA), and isolated MTEC using enzymatic digestion, as previously described.^44,50,51^ We suspended MTEC in a defined proliferation media [base media with 10 μg/mL insulin, 5 μg/mL apo-transferrin, 0.1 μg/mL cholera toxin, 25 ng/mL epidermal growth factor, 30 μg/mL bovine pituitary extract, and 50 nM retinoic acid (Sigma-Aldrich, St. Louis, MO, USA)] to culture at a density of 1.5×104 cells/well in collagen-coated 24-Transwell^®^ plates (Corning, Corning, NY, USA). We allowed organotypic cultures to proliferate with media in both apical and basal compartments starting on day *in vitro* (DIV) 0. We changed the defined proliferation media every other day until DIV 7-9, when transepithelial electrical resistance (TEER) exceeded 1000 Ω×cm2, which marked the end of proliferation.

For the “Normal” phenotype, we allowed organotypic cultures to differentiate at an air-liquid interface (ALI) in a defined differentiation media [base media with 2% v/v NuSerum (BD BioSciences, San Jose, CA, USA), and 50 nM retinoic acid (Sigma-Aldrich, St. Louis, MO, USA)] in the basal compartment starting on DIV 7. We changed the defined differentiation media every other day until DIV 28, which marked the end of differentiation.

For the “T2-Skewed” phenotype, we allowed organotypic cultures to differentiate at an ALI in a defined differentiation media supplemented with IL-13 [base media with 2% v/v NuSerum (BD BioSciences, San Jose, CA, USA), 50 nM retinoic acid (Sigma-Aldrich, St. Louis, MO, USA)], and 25 ng/mL IL-13 (PeproTech, Rocky Hill, NJ, USA)]. We identified this concentration of IL-13 in a preliminary study and supplemented it to the defined proliferation media starting on DIV 5, and then to the defined differentiation media from DIV 7-28, which we changed every other day until DIV 28.

### Immunohistochemistry

We used immunohistochemistry (IHC) to characterize organotypic morphology in unexposed organotypic cultures on DIV 28. We washed organotypic cultures three times with 1× PBS (each wash for 10 min), fixed in 4% PFA (for 24 h at 4°C), embedded in paraffin, sectioned, and then stained with hematoxylin and eosin using an Autostainer XL (Leica Biosystems, Buffalo Grove, IL, USA). We acquired images at 40× on an Eclipse 90i light microscope and processed using associated digital microscopy software (Nikon Instruments, Melville, NY, USA).

### Quantitative reverse transcription-polymerase chain reaction

We used quantitative reverse transcription polymerase chain reaction (qRT-PCR) to quantify mRNA expression for markers of barrier function, cell populations, allergic responses (airway remodeling and hyperresponsiveness), as well as type 1 (T1), T2, and pro-T2 cytokines in unexposed organotypic cultures on DIV 28. We isolated total RNA from organotypic cultures using a RNEasy Micro Kit (Qiagen, Hilden, Germany), and synthesized cDNA using a RevertAid RT Reverse Transcription Kit (ThermoFisher Scientific, Waltham, MA, USA). We performed qRT-PCR using 20 ng cDNA per reaction with commercially available primer probe sets (Table 1), and master mix (Integrated DNA Technologies, Coralville, IA, USA) on an Applied Biosystems^®^ 7900HT Fast Real-Time PCR System (Applied BioSystems, Foster City, CA, USA). We normalized data to a baseline condition of the Ct values for each gene averaged across AJ:Normal, B6:Normal, AJ:T2-Skewed, and B6:T2-Skewed and then to the average Ct values of housekeeping genes Hprt1, Pol2ra, and Tbp using the 2–ΔΔCt method, as previously described.^43^

### AgNP exposure

We exposed organotypic cultures in the apical compartment on DIV 28 to either 2 mM sodium citrate (vehicle control; 0 μg AgNP/mL media), or to silver nanoparticles (AgNP; 20 nm, gold-core, citrate-coated, at 1 mg/mL in 2 mM sodium citrate; nanoComposix, San Diego, CA, USA) for an acute exposure of 24 hours, or a subacute exposure of 4 hours, every other day, over 5 days (5×4 h) at nominal doses of 12.5, 25, or 50 μg AgNP/mL media, as freshly prepared in suspensions of defined differentiation media.

### Inductively coupled plasma-mass spectrometry

We used inductively coupled plasma mass spectrometry (ICP-MS) to quantify dosimetric doses of silver (Ag), gold (Au), and the ratio of silver to gold (Ag:Au) mass associated on exposed organotypic cultures. We washed organotypic cultures three times, each wash for 10 m, with 1× PBS to remove unassociated AgNP. We lysed organotypic cultures with CelLytic M Cell Lysis Reagent (Sigma-Aldrich, St. Louis, MO, USA) for 10 m at 4°C and pelleted the lysates at 50,000×g for 1 h at 4°C using an Optima MAX-XP ultracentrifuge (Beckman Coulter, Brea, CA, USA). We resuspended cell fractions in 500 μL water and stored the samples at 4°C until analysis, as previously described.^41^ We used a Bradford assay (Bio-Rad Laboratories, Hercules, CA, USA) to measure protein content, as previously described.^52^ We normalized data by dividing the Ag, Au, and Ag:Au mass in each cell fraction by the protein content. The Ag:Au mass is an indirect measurement of the colocation of Ag and Au mass, and a high Ag:Au mass therefore suggests that the two metals have not separated through dissolution.

### Transepithelial electrical resistance assay

We used a EVOM2 Epithelial Volt/Ohm Meter (World Precision Instruments, Sarasota, FL, USA) to quantify TEER as a measure of barrier function in exposed organotypic cultures, as previously described.^53^ We normalized data by subtracting the average background resistance (in 1× PBS) and multiplying by the area of the semi-permeable transwell membrane (0.33 cm2).

### 2,3-Naphthalenedicarboxaldehyde assay

We used a 2,3-naphthalenedicarboxaldehyde (NDA) assay to quantify NDA fluorescence at 472/528 nm as a measure of GSH depletion/oxidative stress in exposed organotypic cultures, as previously described.^54^ We normalized data by subtracting the average background absorbance of defined differentiation media, multiplying by a sample dilution factor (1/100) for fitting to a standard curve of known GSH concentrations in media, and dividing by the protein content.

### 2,7-Dichlorofluorescein assay

We used a 2,7-dichlorofluorescein (DCF) assay (Abcam, Cambridge, MA, USA) to quantify DCF fluorescence at 485/535 nm as a measure of ROS production/oxidative stress in exposed organotypic cultures, as previously described.^55^ We normalized data by subtracting the average background absorbance of defined differentiation media and dividing by the DCF fluorescence of vehicle controls.

### Malondialdehyde assay

We used a malondialdehyde (MDA) assay (Abcam, Cambridge, MA, USA) to quantify MDA fluorescence at 523/553 nm as a measure of lipid peroxidation/oxidative stress in exposed organotypic cultures, as previously described.^56^ We normalized data by subtracting the average background absorbance in media, multiplying by a correction factor (4) for using 200 μL of the 800 μL reaction mix, multiplying by a dilution factor (1/100) for fitting to a standard curve of known MDA concentrations in media, and dividing by the protein content.

### Lactate dehydrogenase assay

We used a lactate dehydrogenase (LDH) assay (Promega, Madison, WI, USA) to quantify LDH absorbance in cell culture supernatant at 490 nm as a measure of cytotoxicity in exposed organotypic cultures, as previously described.^57^ We normalized data by subtracting the average background absorbance in media and dividing by the LDH absorbance of positive controls.

### Statistics

We used linear mixed effects models to test the fixed effects of genotype, phenotype, and exposure using R statistical software.^58^ We adjusted each model for random effects to account for variability across and within biological and technical replicates. We used ANOVA to test the significance of each fixed effect as well as interactions between these fixed effects using the nominal dose of AgNP mass as a categorical variable. We tested the significance of interactions between these fixed effects to characterize G×P effects on AgNP toxicity. For all analyses, we considered an effect with a P-value less than 0.05 (P<0.05) statistically significant and adjusted all effects for multiple comparisons when comparing each nominal dose of AgNP mass to its vehicle control. We used a BMD approach to characterize G×P effects on the sensitivity of adverse cellular responses by accounting for differences in the effective dose ranges to induce AgNP toxicity measured by the nominal and dosimetric BMD and their lower 95% confidence limits (BMDL) across genotypes, phenotypes, and exposures using the dosimetric dose of Ag mass as a continuous variable **(Equation 1)**.

## Results

### The “T2-skewed” phenotype is an in vitro model of chronic respiratory diseases

We compared genotype and phenotype effects for organotypic morphology and mRNA expression to achieve baseline characterization of organotypic cultures. We did not observe genotype effects on organotypic morphology; however, we observed phenotype effects on organotypic morphology **(Figure 1)**. We observed organotypic morphology of the “T2-Skewed” phenotype recapitulated clinical features of airway remodeling, marked by a shift toward a pseudostratified columnar epithelium with an abundance of epithelial glycoproteins. We observed an increase in epithelial thickness due to cell proliferation (marked by blue-stained nuclei), shifted the simple cuboidal epithelium under the “Normal” phenotype to the pseudostratified columnar epithelium under the “T2-Skewed” phenotype. An increase in the density of ciliated cells is a distinguishing feature of pseudostratified columnar epithelium, which we also observed on the apical surface under the “T2-Skewed” phenotype. An increase in epithelial glycoproteins, including mucus and chitin (marked by unstained or light pink-stained inclusion bodies), is a distinguishing feature of T2 airway inflammation, which we also observed under the “T2-Skewed” phenotype.

**Figure 1.**
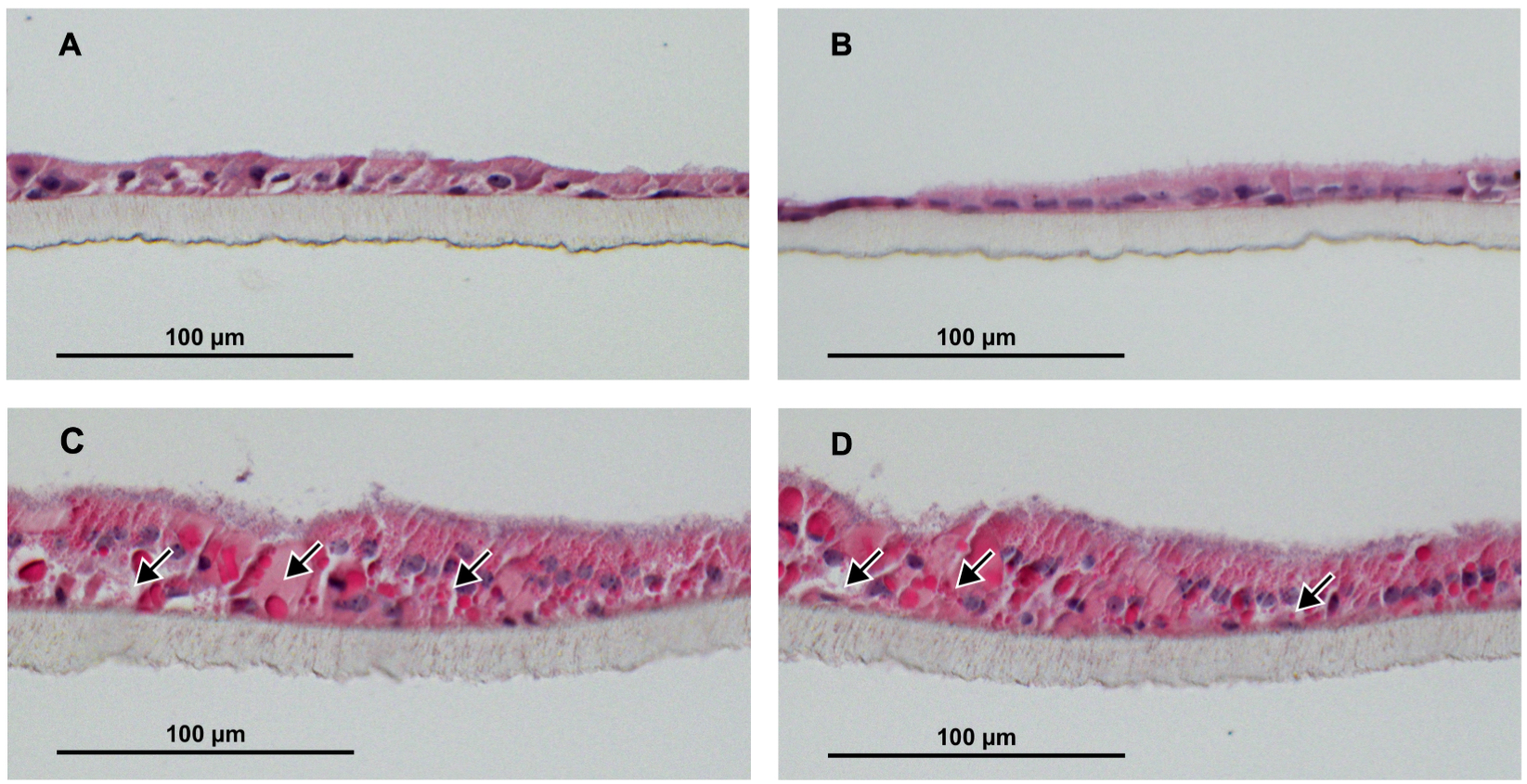
The “T2-Skewed” phenotype is characterized by organotypic morphology associated with clinical features of chronic respiratory diseases, or a pseudostratified columnar epithelium with an abundance of epithelial glycoproteins at baseline (DIV 28). Representative H&E staining of untreated organotypic cultures **(A)** AJ:Normal, **(B)** B6:Normal, **(C)** AJ:T2-Skewed, and **(D)** B6:T2-Skewed on DIV 28. Scale bar: 100 μm. Arrows: epithelial glycoproteins.

We did not observe genotype effects on mRNA expression (*P*>0.05); however, we observed phenotype effects on mRNA expression (*P*<0.001) **(Figure 2; Supplementary Data, Table S1)**. We observed mRNA expression under the “T2-Skewed” phenotype recapitulated clinical features of chronic respiratory diseases, including barrier dysfunction,^12^ mucus production,^9^ allergic responses,^10^ and T2 responses.^47,59^ We observed barrier dysfunction under the “T2-Skewed” phenotype by downregulation of *Tuba1a*, *Tubb4a*, *Tjp1*, *Jup*, and *Gja1*, which are markers for α-tubulin, β-tubulin, tight, adherens, and gap junctions, respectively. We observed genotype effects on mRNA expression for *Tubb4a*, *Jup*, and *Gja1* under the “Normal” phenotype (*P*<0.001 - *P*<0.05). We observed mucus production under the “T2-Skewed” phenotype by upregulation of *Muc5ac*, a marker for mucin cells; however, we did not observe mucin cell metaplasia under the “T2-Skewed” phenotype due to no evidence of reciprocal regulation with *Foxj1*, a marker for ciliated cells. We observed genotype effects on mRNA expression for *Scgb1a1* and *Muc5ac* under the “T2-Skewed” phenotype (*P*<0.001 - *P*<0.01). We observed allergic responses associated with airway remodeling and hyperresponsiveness under the “T2-Skewed” phenotype by upregulation of *Pcna*, *Ym1/2*, and *Pla2g10*, which are markers for cellular proliferation, agglutination, and eicosanoid production, respectively. We observed genotype effects on mRNA expression for *Pcna* and *Pla2g10* under the “T2-Skewed” phenotype (*P*<0.001). We observed reciprocal regulation of T1/T2 responses under the “T2-Skewed” phenotype by downregulation of T1 cytokines *Il2*, *Infg*, and *Tnfb* and upregulation of upstream pro-T2 cytokines *Il25*, *Il33*, and *Tslp* as well as downstream T2 cytokines *Il4*, *Il5*, *Il6*, *Il9*, *Il10*, and *Il13*. We observed genotype effects on mRNA expression for *Il2* and *Infg* under the “Normal” phenotype (*P*<0.001) as well as *Tnfb*, *Il33*, *Il6*, *and Il10* under the “T2-Skewed” phenotype (*P*<0.001 - *P*<0.01).

**Figure 2.**
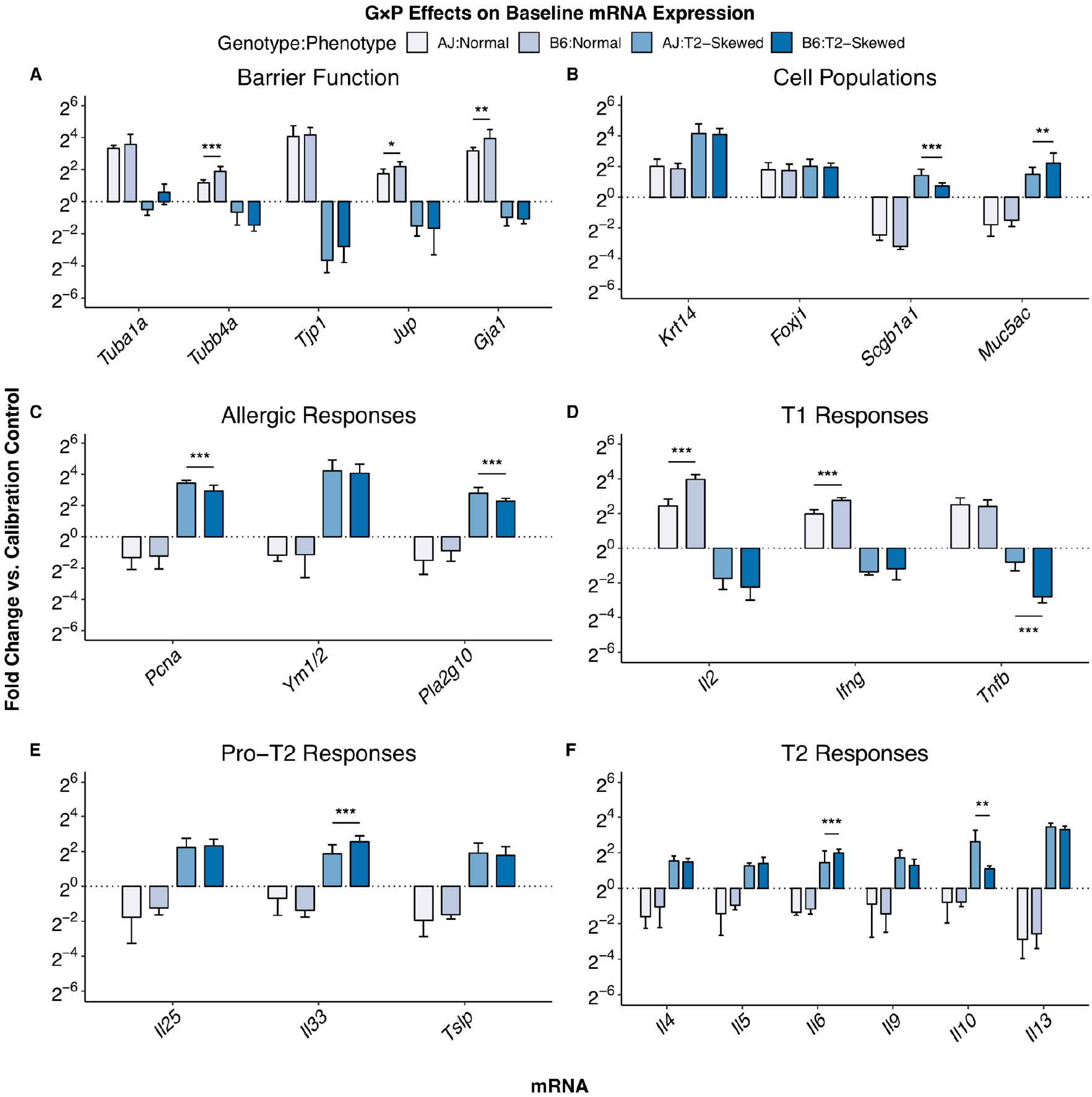
The “T2-Skewed” phenotype is characterized by mRNA expression associated with clinical features of chronic respiratory diseases, including barrier dysfunction, mucus production, allergic responses, and T2 responses at baseline (DIV 28). Total RNA was isolated from untreated organotypic cultures on DIV 28, and cDNA was subjected to qRT-PCR analysis for mRNA expression of markers associated with **(A)** barrier function, **(B)** cell populations, **(C)** allergic responses, **(D)** T1 responses, **(E)** pro-T2 responses, and **(F)** T2 responses. The fold change in mRNA expression was compared between genotypes (asterisks) for each phenotype. *n* = 3 biological replicates with technical replicates. Data represent mixed effects estimate ± 95% CI; NS (*P*>0.05); * (*P*<0.05); *** (*P*<0.001).

### Exposure and phenotype effects on the linear relationship between the nominal dose of AgNP mass and the dosimetric dose of Ag, Au, and Ag:Au mass

We compared genotype, phenotype, and exposure effects for linear relationships between the nominal dose of AgNP mass and the dosimetric dose of Ag, Au, and Ag:Au mass. We observed positive linear relationships between the nominal dose of AgNP mass and the dosimetric dose of Ag **(Figure 3A-B; Supplementary Data, Table S2-S3)** and Au mass **(Figure 3C-D; Supplementary Data, Table S2-S3)** for AJ:Normal, B6:Normal, AJ:T2-Skewed, and B6:T2-Skewed at the acute 24 h exposure and the subacute 5×4 h exposure. We did not observe genotype or phenotype effects on the dosimetric dose of Ag and Au mass (*P*>0.05); however, we observed exposure effects on the dosimetric dose of Ag and Au mass, with the highest dosimetric dose of Ag and Au mass detected at the acute 24 h exposure (*P*<0.001). Additionally, we observed G×P effects on the dosimetric dose of Ag and Au mass at the acute 24 h exposure and the subacute 5×4 h exposure (*P*<0.001). We did not observe a linear relationship between the nominal dose of AgNP mass and the dosimetric dose of Ag:Au mass **(Figure 3E-F; Supplementary Data, Table S2-S3)** for AJ:Normal, B6:Normal, AJ:T2-Skewed, and B6:T2-Skewed at the acute 24 h exposure and the subacute 5×4 h exposure. We did not observe genotype or exposure effects on the dosimetric dose of Ag:Au mass (*P*>0.05); however, we observed phenotype effects on the dosimetric dose of Ag:Au mass, with the lowest dosimetric dose of Ag:Au mass detected in the “T2-Skewed” phenotype (*P*<0.05). Additionally, we observed G×P effects on the dosimetric dose of Ag:Au mass at the acute 24 h exposure (*P*<0.05) and the subacute 5×4 h exposure (*P*<0.001).

**Figure 3.**
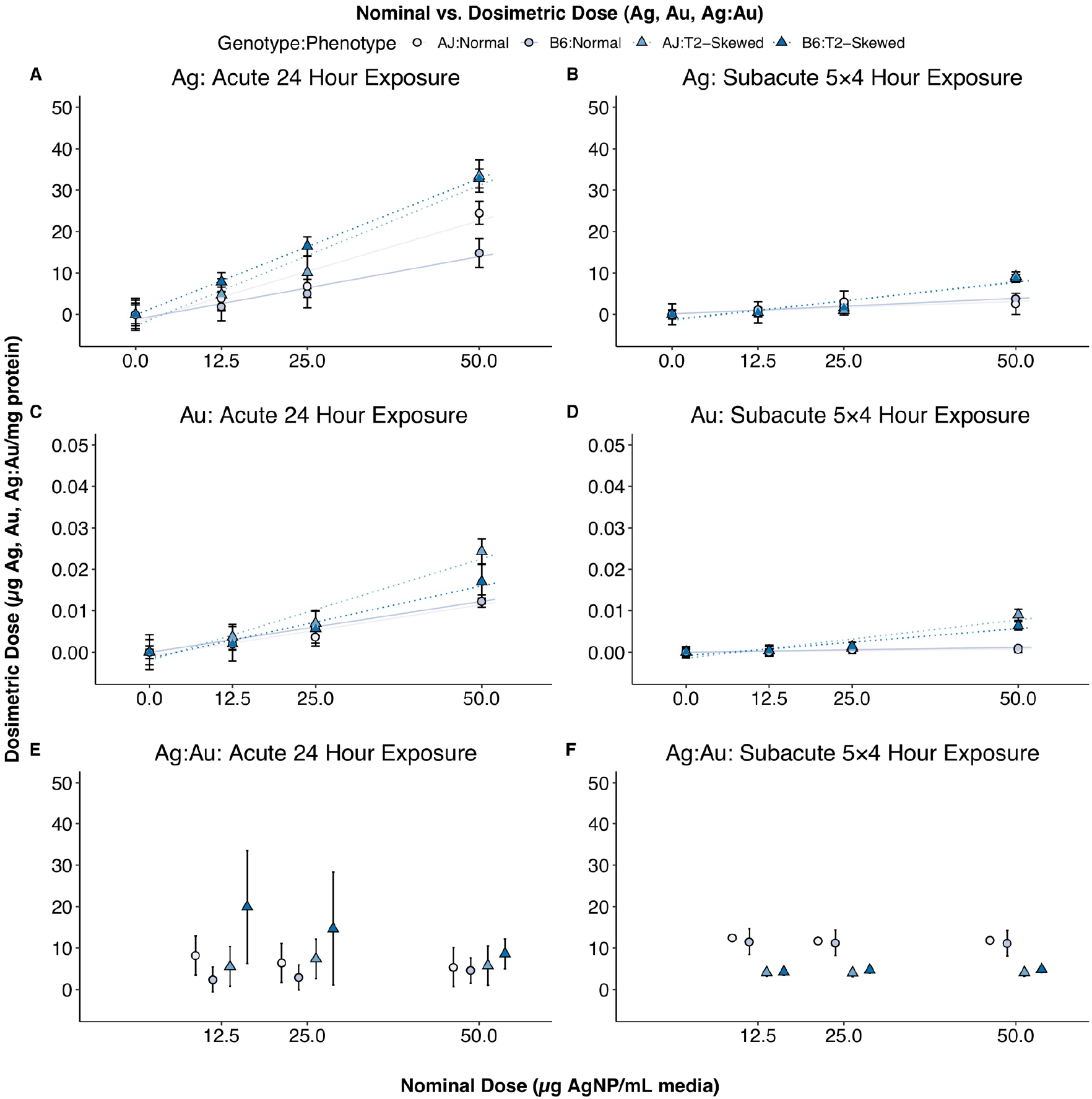
Phenotype and exposure effects on the linear relationship between the nominal dose of AgNP mass and the dosimetric doses of Ag, Au, and Ag:Au mass. Organotypic cultures were exposed to nominal doses of 0, 12.5, 25, or 50 μg AgNP/mL media at an acute 24 h exposure and a subacute 5×4 h exposure, and cell lysates were subjected to ICP-MS analysis for dosimetric doses of Ag **(A, B)**, Au **(C, D)**, and Ag:Au **(E, F)** mass. Dosimetric doses of Ag and Au mass were normalized to the protein content and were then divided to derive dosimetric doses of Ag:Au mass. *n* = 3 biological replicates. Data represent mixed effects estimate ± 95% CI.

### Genotype and phenotype effects on AgNP-induced barrier dysfunction

We compared genotype, phenotype, and exposure effects for nominal and dosimetric dose-response relationships for TEER as a measure of barrier function. We observed negative dose-response relationships for AgNP-induced barrier dysfunction for AJ:T2-Skewed at the acute 24 h exposure and the subacute 5×4 h exposure (*P*<0.001) **(Figure 4; Supplementary Data, Tables S4-S5)**. We observed genotype and phenotype effects on AgNP-induced barrier dysfunction (*P*<0.001), with increased sensitivity to barrier dysfunction detected in AJ:T2-Skewed. We did not observe exposure effects on AgNP-induced barrier dysfunction (*P*>0.05); however, we observed G×P effects on AgNP-induced barrier dysfunction at the acute 24 h exposure and the subacute 5×4 h exposure (*P*<0.001).

**Figure 4.**
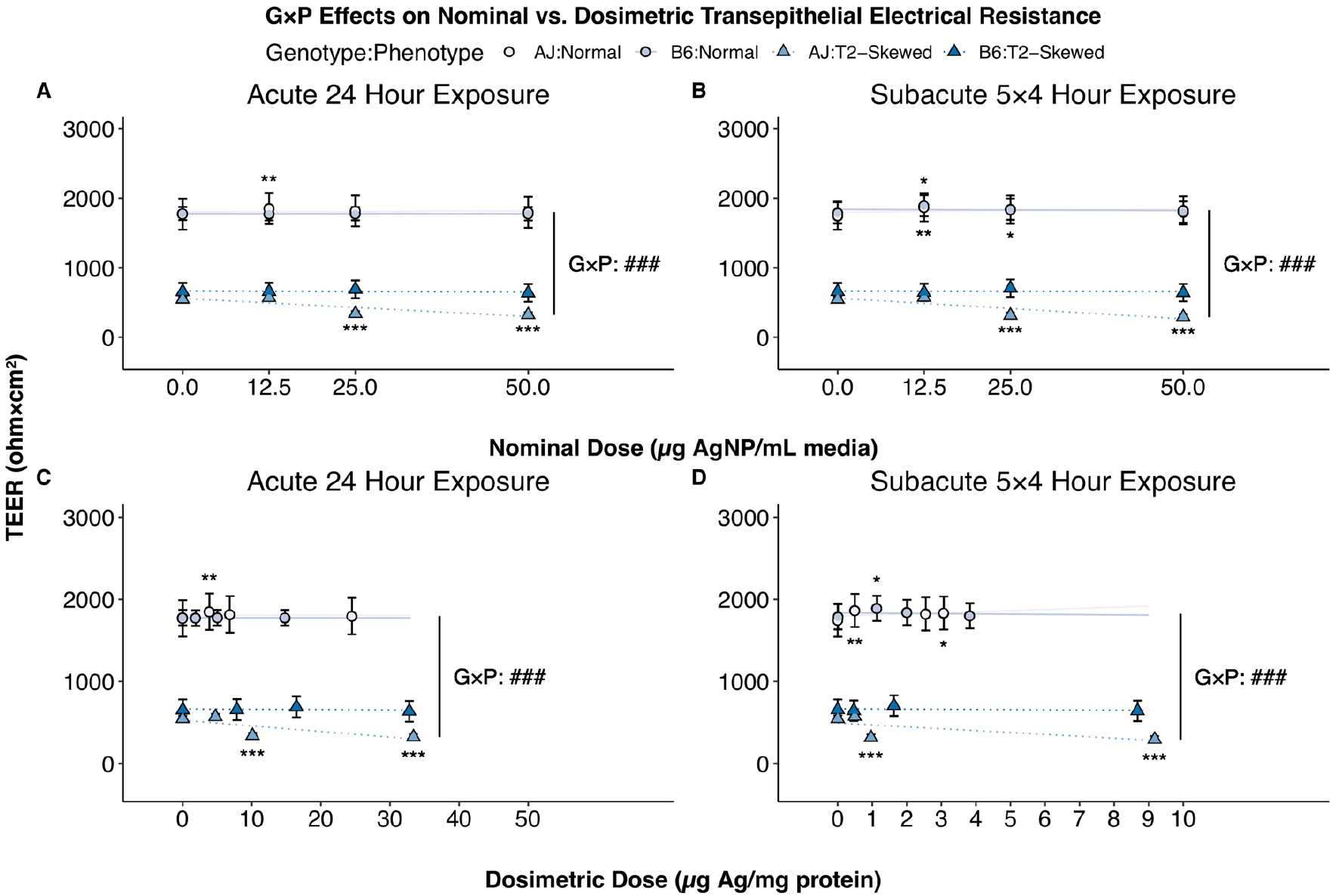
Genotype and phenotype effects on AgNP-induced barrier dysfunction. TEER, a non-specific method to assay barrier function in epithelial monolayers, was assayed in organotypic cultures exposed to nominal doses of 0, 12.5, 25, or 50 μg AgNP/mL media at an acute 24 h exposure and a subacute 5×4 h exposure. TEER was normalized to the background resistance and area of the semi-permeable transwell membrane, then compared to vehicle controls (asterisks) for each genotype and phenotype; G×P effects were compared across genotypes and phenotypes (pounds) for each exposure across nominal **(A, B)** and dosimetric **(C, D)** dose-response relationships. *n* = 3 biological replicates with technical replicates. Data represent mixed effects estimate ± 95% CI; NS (*P*>0.05); * (*P*<0.05); ** (*P*<0.01); *** (*P*<0.001); ### (*P*<0.001).

### Genotype and phenotype effects on AgNP-induced GSH depletion

We compared genotype, phenotype, and exposure effects for nominal and dosimetric dose-response relationships for NDA fluorescence as a measure of GSH content. We observed negative dose-response relationships for AgNP-induced GSH depletion for AJ:Normal, B6:Normal, AJ:T2-Skewed, and B6:T2-Skewed at the acute 24 h exposure and the subacute 5×4 h exposure (*P*<0.001 - *P*<0.05) **(Figure 5; Supplementary Data, Tables S4-S5)**. We observed genotype and phenotype effects on AgNP-induced GSH depletion (*P*<0.001), with increased sensitivity to GSH depletion detected in AJ:T2-Skewed. We did not observe exposure effects on AgNP-induced GSH depletion (*P*>0.05); however, we observed G×P effects on AgNP-induced GSH depletion at the acute 24 h exposure and the subacute 5×4 h exposure (*P*<0.001).

**Figure 5.**
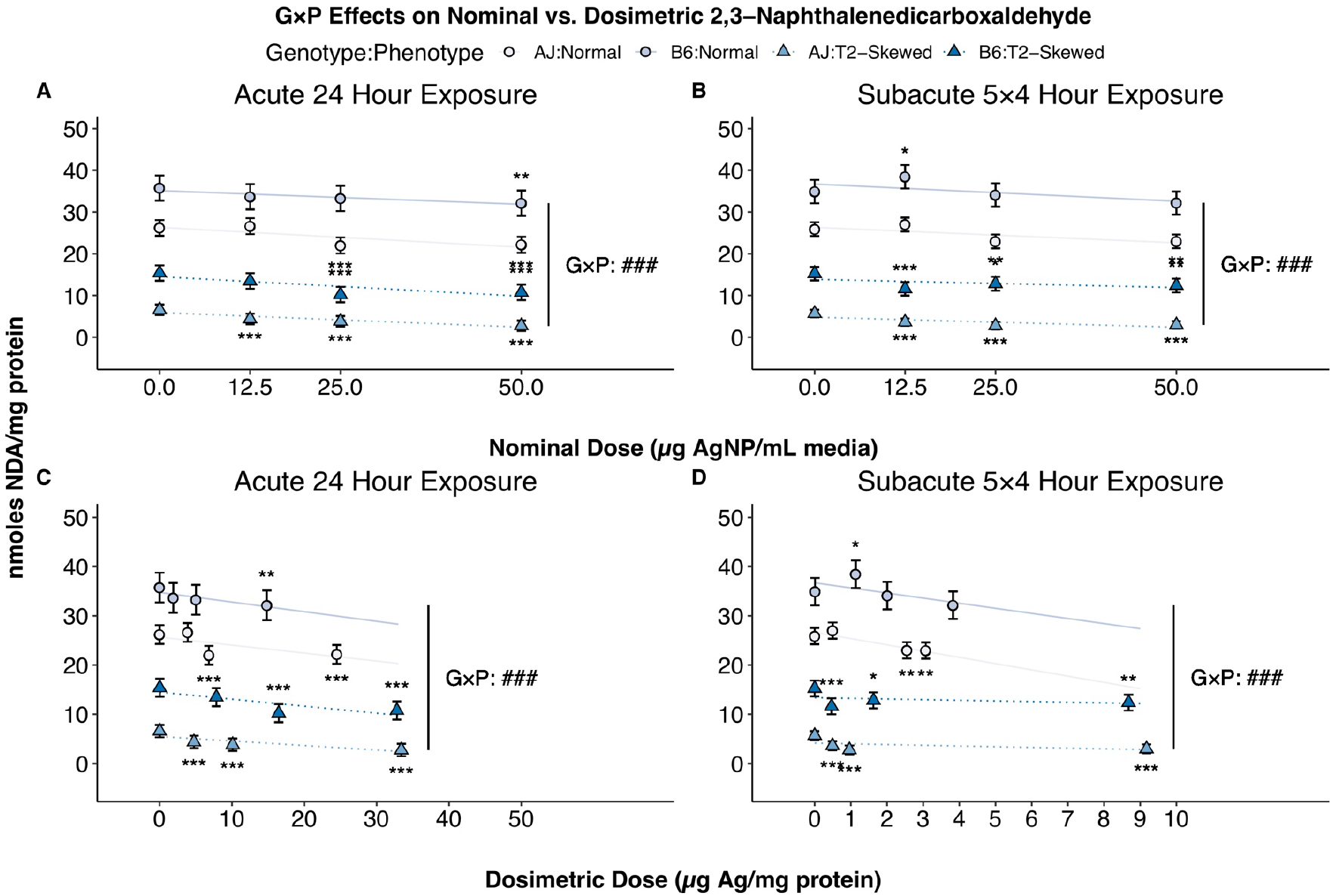
Genotype and phenotype effects on AgNP-induced GSH depletion. NDA fluorescence, a specific method to assay dissolved GSH concentrations in media, was assayed in organotypic cultures exposed to nominal doses of 0, 12.5, 25, or 50 μg AgNP/mL media at an acute 24 h exposure and a subacute 5×4 h exposure. NDA fluorescence was normalized to the protein content, then compared to vehicle controls (asterisks) for each genotype and phenotype; G×P effects were compared across genotypes and phenotypes (pounds) for each exposure across nominal **(A, B)** and dosimetric **(C, D)** dose-response relationships. *n* = 3 biological replicates with technical replicates. Data represent mixed effects estimate ± 95% CI; NS (*P*>0.05); * (*P*<0.05); ** (*P*<0.01); *** (*P*<0.001); ### (*P*<0.001).

### Genotype, phenotype, and exposure effects on AgNP-induced ROS production

We compared genotype, phenotype, and exposure effects for nominal and dosimetric dose-response relationships for DCF fluorescence as a measure of ROS production. We observed positive dose-response relationships for AgNP-induced ROS production for AJ:Normal, B6:Normal, AJ:T2-Skewed, and B6:T2-Skewed at the acute 24 h exposure and the subacute 5×4 h exposure (*P*<0.001) **(Figure 6; Supplementary Data, Tables S4-S5)**. We observed genotype, phenotype, and exposure effects on AgNP-induced ROS production (*P*<0.001), with increased sensitivity to ROS production detected in AJ:T2-Skewed at the acute 24 h exposure. Additionally, we observed G×P effects on AgNP-induced ROS production at the acute 24 h exposure and the subacute 5×4 h exposure (*P*<0.001).

**Figure 6.**
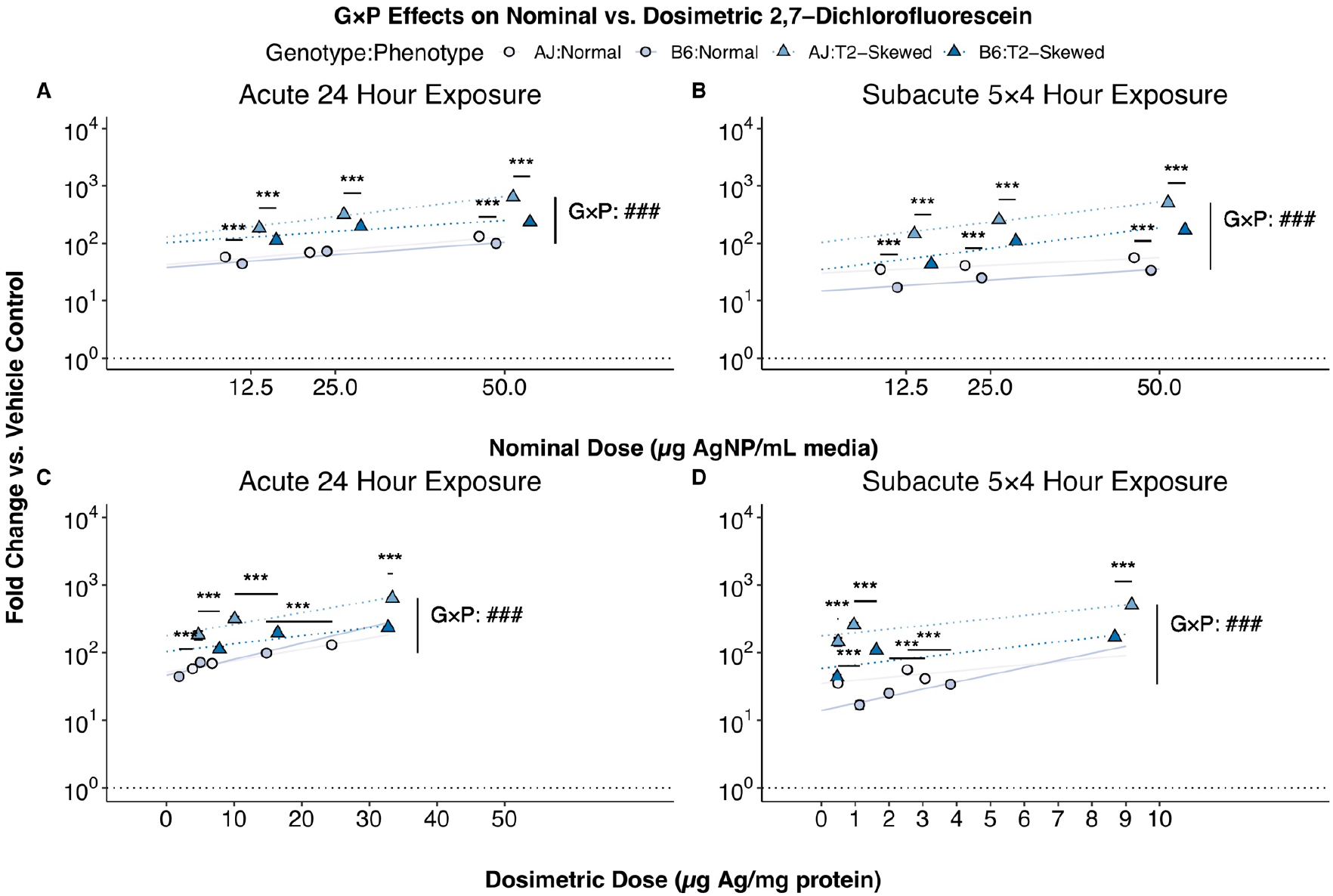
Genotype, phenotype, and exposure effects on AgNP-induced ROS production. DCF fluorescence, a non-specific method to assay ROS production in media, was assayed in organotypic cultures exposed to nominal doses of 0, 12.5, 25, or 50 μg AgNP/mL media at an acute 24 h exposure and a subacute 5×4 h exposure. The fold change in DCF fluorescence was normalized to the vehicle controls, then compared between genotypes (asterisks) for each phenotype; G×P effects were compared across genotypes and phenotypes (pounds) for each exposure across nominal **(A, B)** and dosimetric **(C, D)** dose-response relationships. *n* = 3 biological replicates with technical replicates. Data represent mixed effects estimate ± 95% CI; NS (*P*>0.05); *** (*P*<0.001); ### (*P*<0.001).

### Genotype, phenotype, and exposure effects on AgNP-induced lipid peroxidation

We compared genotype, phenotype, and exposure effects for nominal and dosimetric dose-response relationships for MDA fluorescence as a measure of lipid peroxidation. We observed positive dose-response relationships for AgNP-induced lipid peroxidation for AJ:Normal, B6:Normal, AJ:T2-Skewed, and B6:T2-Skewed at the acute 24 h exposure and the subacute 5×4 h exposure (*P*<0.001) **(Figure 7; Supplementary Data, Tables S4-S5)**. We observed genotype, phenotype, and exposure effects on AgNP-induced lipid peroxidation (*P*<0.001), with increased sensitivity to lipid peroxidation detected in AJ:T2-Skewed at the acute 24 h exposure. Additionally, we observed G×P effects on AgNP-induced lipid peroxidation at the acute 24 h exposure and the subacute 5×4 h exposure (*P*<0.001).

**Figure 7.**
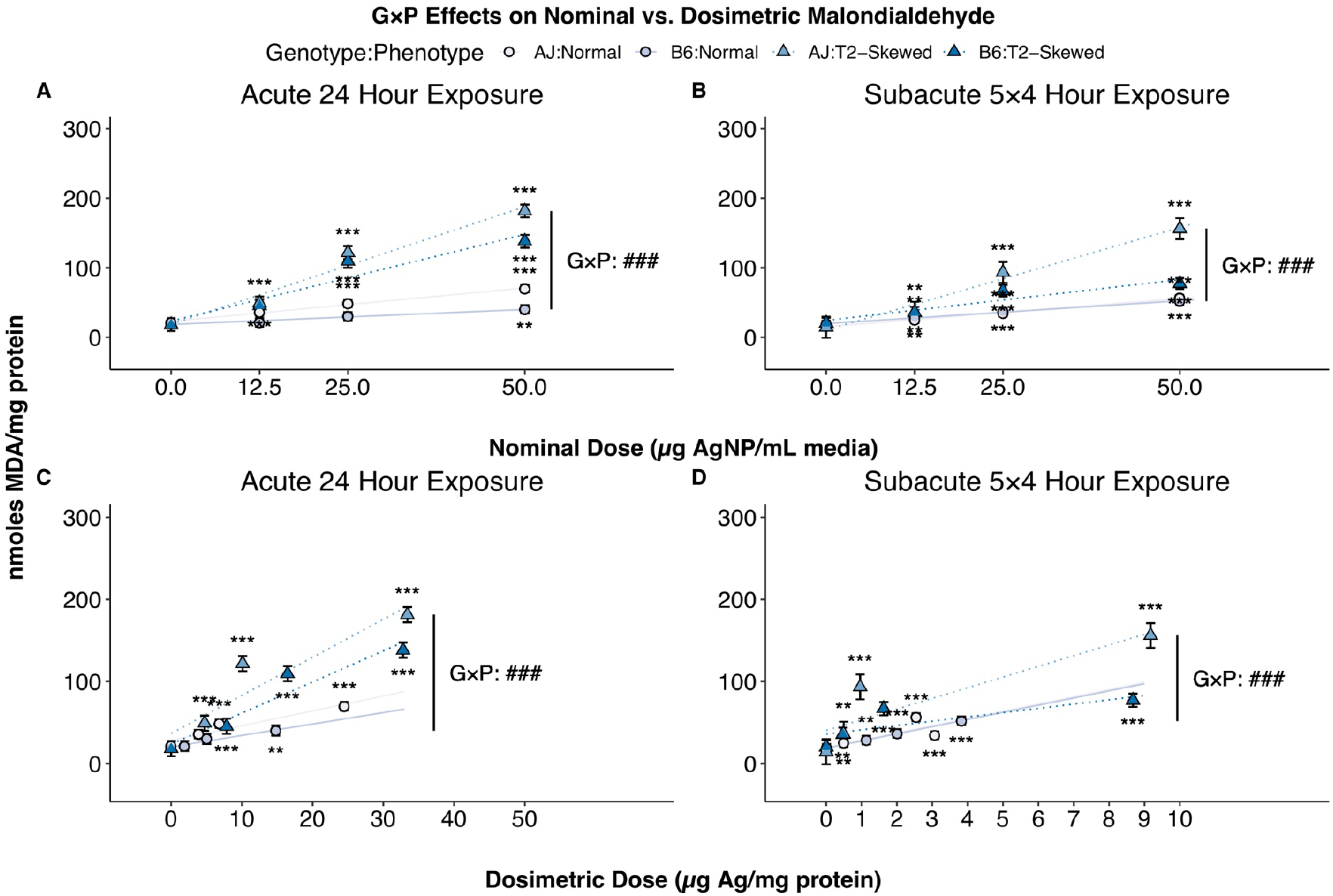
Genotype, phenotype, and exposure effects on AgNP-induced lipid peroxidation. MDA fluorescence, a specific method to assay dissolved MDA concentrations in media, was assayed in organotypic cultures exposed to nominal doses of 0, 12.5, 25, or 50 μg AgNP/mL media at an acute 24 h exposure and a subacute 5×4 h exposure. MDA fluorescence was normalized to the protein content, then compared to vehicle controls (asterisks) for each genotype and phenotype; G×P effects were compared across genotypes and phenotypes (pounds) for each exposure across nominal **(A, B)** and dosimetric **(C, D)** dose-response relationships. *n* = 3 biological replicates with technical replicates. Data represent mixed effects estimate ± 95% CI; NS (*P*>0.05); ** (*P*<0.01); *** (*P*<0.001); ### (*P*<0.001).

### Genotype and phenotype effects on AgNP-induced cytotoxicity

We compared genotype, phenotype, and exposure effects for nominal and dosimetric dose-response relationships for LDH absorbance as a measure of cytotoxicity. We observed positive dose-response relationships for AgNP-induced cytotoxicity for AJ:Normal, B6:Normal, AJ:T2-Skewed, and B6:T2-Skewed at the acute 24 h exposure and the subacute 5×4 h exposure (*P*<0.001) **(Figure 8; Supplementary Data, Tables S4-S5)**. We observed genotype and phenotype effects on AgNP-induced cytotoxicity (*P*<0.001), with increased sensitivity to cytotoxicity detected in AJ:T2-Skewed. We did not observe exposure effects on AgNP-induced cytotoxicity (*P*>0.05); however, we observed G×P effects on AgNP-induced cytotoxicity at the acute 24 h exposure and the subacute 5×4 h exposure (*P*<0.001).

**Figure 8.**
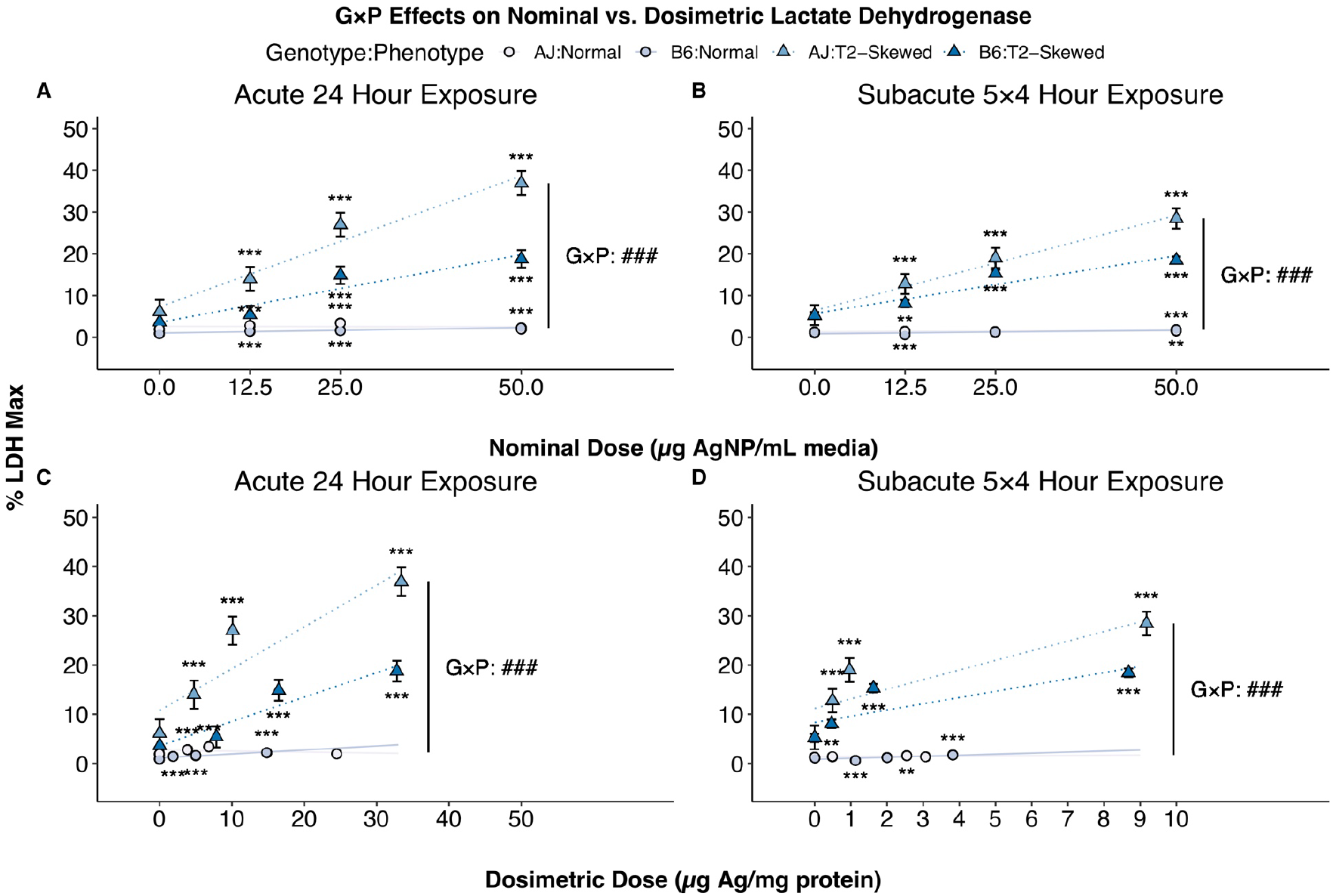
Genotype and phenotype effects on AgNP-induced cytotoxicity. LDH absorbance, a specific method to assay dissolved LDH concentrations in media, was assayed in organotypic cultures exposed to nominal doses of 0, 12.5, 25, or 50 μg AgNP/mL media at an acute 24 h exposure and a subacute 5×4 h exposure. LDH absorbance was normalized to positive controls, then compared to vehicle controls (asterisks) for each genotype and phenotype; G×P effects were compared across genotypes and phenotypes (pounds) for each exposure across nominal **(A, B)** and dosimetric **(C, D)** dose-response relationships. *n* = 3 biological replicates with technical replicates. Data represent mixed effects estimate ± 95% CI; NS (*P*>0.05); * (*P*<0.05); ** (*P*<0.01); *** (*P*<0.001); ### (*P*<0.001).

### Genotype, phenotype, and exposure effects on sensitivity to AgNP-induced adverse cellular responses

We compared genotype, phenotype, and exposure effects for nominal and dosimetric BMD (BMDL) as a measure of sensitivity to adverse cellular responses by accounting for differences in the effective dose ranges to induce AgNP toxicity. We observed similar patterns of nominal and dosimetric BMD (BMDL) across genotypes, phenotypes, and exposures for each adverse cellular response **(Figure 9; Supplementary Data, Table S6)**. The most sensitive adverse cellular response across nominal and dosimetric dose-response relationships was marked by the lowest effective dose to induce AgNP toxicity, which was ROS production at the acute 24 h exposure and the subacute 5×4 h exposure.

**Figure 9.**
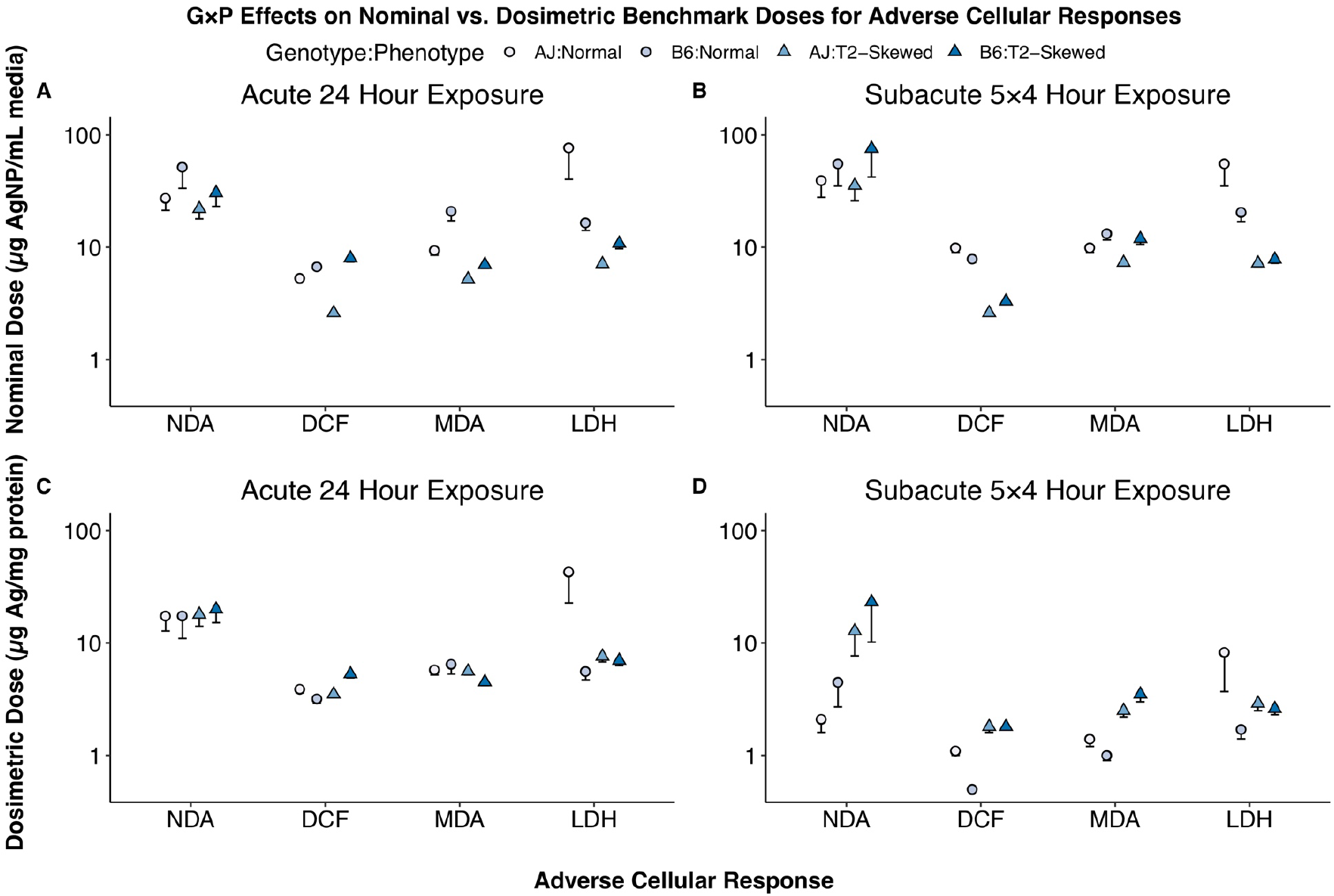
Genotype, phenotype, and exposure effects on sensitivity to AgNP-induced adverse cellular responses. BMD (BMDL) were used to compare sensitivity to adverse cellular responses. BMD (BMDL) were derived from significant nominal **(A, B)** and dosimetric **(C, D)** dose-response relationships for each genotype, phenotype, and exposure using the dosimetric dose of Ag mass as a continuous variable. *n* = 3 biological replicates with technical replicates. Data represent BMD - BMDL.

## Discussion

This is the first study to use organotypic cultures as a high-content *in vitro* model of the conducting airway to characterize G×P effects on AgNP toxicity. We characterized nominal and dosimetric dose-response relationships for AgNP-induced barrier dysfunction, GSH depletion, ROS production, lipid peroxidation, and cytotoxicity across genotypes, phenotypes, and exposures to understand G×P effects on AgNP toxicity. Across each of the nominal and dosimetric dose-response relationships, we observed increased sensitivity to AgNP toxicity under AJ:T2-Skewed at the acute 24 h exposure. We observed higher dosimetric doses of Ag and Au mass under the “T2-Skewed” phenotype at the acute 24 h exposure and lower dosimetric doses of Ag:Au mass under the “T2-Skewed” phenotype at the subacute 5×4 h exposure, suggesting that phenotype and exposure effects on association and dissolution are greatest for the “T2-Skewed” phenotype at acute and subacute exposures, respectively. The dosimetric dose of Ag:Au mass is an indirect measurement of the colocation of Ag and Au mass, and a low Ag:Au mass suggests that the two metals have separated through dissolution, potentially leading to increased AgNP bioavailability and bioactivity. We used a BMD approach to characterize G×P effects on the sensitivity of adverse cellular responses by accounting for differences in the effective dose ranges to induce AgNP toxicity across genotypes, phenotypes, and exposures, and identified ROS production as the most sensitive adverse cellular response at the acute 24 h exposure and the subacute 5×4 h exposure.

These data suggest that phenotype and exposure effects on AgNP particokinetics are greatest for the “T2-Skewed” phenotype at the acute 24 h exposure, rather than the subacute 5×4 h exposure. For the subacute 5×4 h exposure, when the delivered dose was separated across days, the peak exposures per day seemed to drive the overall exposure. The subacute 5×4 h exposure was chosen to model a work-week exposure scenario, as previous studies have suggested a potential high-risk for AgNP exposures in occupational settings.^21,60–63^ Together, this highlights the importance of using multiple genotypes, phenotypes, and exposures when screening and prioritizing other potential respiratory toxicants in order to produce more biologically robust predictions of adverse organism responses resulting from occupational exposures.^64^

We used dosimetric dose-response relationships to account for genotype, phenotype, and exposure effects on AgNP particokinetics and/or airway physiology of organotypic cultures. However, when we used these data to derive dosimetric BMD (BMDL), the sensitivity of the “T2-Skewed” phenotype decreased for the acute 24 h exposure but increased (alongside the “Normal” phenotype) for the subacute 5×4 h exposure. This suggests the importance of considering exposure effects on AgNP toxicity. While electron microscopy may better inform whether these genotype, phenotype, and exposure effects on association and dissolution lead to differences in uptake of whole particles, the approach used in the present study allowed for the detection of AgNP toxicity that can be translated to physiologically-relevant units measurable in occupational settings.

While we addressed some of these differences in AgNP particokinetics and/or airway physiology using dosimetric dose-response relationships, the *in vitro* to *in vivo* extrapolation (IVIVE) from organotypic cultures to the lungs may still exhibit significant variability. This was highlighted by Bachler et al., who developed a physiologically-based pharmacokinetic/toxicokinetic (PBPK) model for AgNP toxicity in various organ systems. For the lungs, the authors calibrated their human respiratory tract model to estimate the deposition fraction of inhaled particles from 1 to 1000 nm among the extrathoracic, endothoracic, bronchial, bronchiolar, and alveolar-interstitial compartments, and accounted for mucociliary clearance rates as well as time-dependent uptake of particles to the systemic blood circulation.^65^ While deposition and dissolution can occur throughout the conducting and respiratory airways, deposition in the conducting airway of healthy individuals will be reduced by mucociliary clearance, with a fraction of deposited Ag mass undergoing dissolution upon inhalation. However, this host defense mechanism may be impaired in individuals with pre-existing chronic respiratory diseases, and raises the potential for increased deposition, dissolution, and thus AgNP toxicity. This could be evaluated in future studies using healthy and allergic mice that also incorporate IVIVE or PBPK models to derive human equivalent concentrations (HEC). Weldon et al. used HEC based upon the dosimetric dose of Ag mass in the alveolar region, which showed the highest retention time and burden compared to other organs as reported in the *in vivo* study by Sung et al., to derive a health-based occupational exposure limit (OEL) of 0.19 μg AgNP/m^3^.^64^ Together, this highlights the importance of considering PBPK parameters within sensitive populations when deriving dose-response relationships, HEC, and OEL which can be used to contribute relevant data and efficient characterization toward risk assessment for occupational exposures.

Our observations were supported by previous *in vitro* and *in vivo* studies on the effects of host genetic and acquired factors on engineered nanomaterial (ENM) toxicity. With regard to host genetic factors, Weldon et al. exposed organotypic cultures derived from primary murine embryonic midbrain cells, and observed size, coating, genotype, and developmental stage effects on AgNP-induced cytotoxicity, with A/J and C57BL/6J mice being differentially sensitive genotypes.^57^ Scoville et al. exposed eight genotypes of male mice and observed genotype effects on quantum dot (QD)-induced T2 lung inflammation, with A/J and C57BL/6J mice being the most differentially sensitive genotypes. The authors quantified dosimetric doses of cadmium (Cd) mass in the lungs of these differentially sensitive genotypes and detected higher burdens in A/J mice compared to C57BL/6J mice, suggesting that airway physiology (marked by alveolar size and airway branching) may be a contributing factor in genotype effects on QD toxicity—which may be broadly applicable to other types of ENM, such as AgNP.^42,66,67^ With regard to host acquired factors, Chuang et al. treated OVA-sensitized, female BALB/cJ mice by inhalation and observed antigen effects on AgNP-induced oxidative stress in allergic mice compared to healthy mice and fresh air controls, with allergic and healthy mice being differentially sensitive to AgNP toxicity.^68^ In a similar study, Alessandrini et al. exposed OVA-sensitized, female BALB/cJ mice by tracheal instillation and observed size, coating, dose, and antigen effects on AgNP-induced T2 lung inflammation in allergic mice compared to healthy mice and vehicle controls, with allergic and healthy mice also being differentially sensitive to AgNP toxicity.^69^ Interestingly, the authors observed AgNP-induced T2 lung inflammation in a biphasic manner, with AgNP_CIT200_ and AgNP_PVP200_ attenuating T2 lung inflammation, and AgNP_CIT50_ and AgNP_PVP50_ exacerbating T2 lung inflammation. In the present study; we observed genotype effects to be more understated than phenotype effects on baseline characterization of organotypic cultures. However, we observed G×P effects on AgNP toxicity and suggest these were driven by phenotype rather than genotype effects on AgNP toxicity.

The “T2-Skewed” phenotype’s sensitivity to AgNP toxicity may be attributed to increased association and dissolution, which can be mediated by particle interactions with chloride ions (Cl^-^) within secreted mucus. Yasuo et al. observed IL-13 induced upregulation of *Muc5ac* and calcium-activated chloride channel 1 (*Clca1*) when skewing differentiation of airway epithelial cells.^70^ Therefore, under the “T2-Skewed” phenotype, IL-13 may induce upregulation of *Clca1* in addition to *Muc5ac*, potentially leading to increased AgNP bioavailability and bioactivity. The “T2-Skewed” phenotype’s sensitivity to AgNP toxicity may also be attributed to impaired ion transport, which can be mediated by amiloride-sensitive epithelial Na^+^ channel (ENaC) activity. Kimura et al. observed E3 ubiquitin-protein ligase NEDD4-like (*Nedd4l*) regulates ENaC activity in the lungs, and under normal conditions, it inhibits ENaC activity to prevent T2 airway inflammation.^71^ Campbell et al. observed a positive association between a 6 kbp deletion in an intron of *Nedd4l* and the odds of asthma (Odds Ratio [OR] = 3.13; *P*<0.05);^72^ and as a critical mediator of asthma, IL-13 may induce downregulation of *Nedd4l* to promote ENaC activity and T2 airway inflammation. The “T2-Skewed” phenotype’s sensitivity to AgNP toxicity may also be attributed to GSH depletion, which can be mediated by T2 responses and/or NF-κB activation. Nam et al. observed IL-13 induced NADPH oxidase activity and iNOS to promote ROS/RNS production and GSH depletion in hippocampal neurons.^73^ Although no studies have confirmed this mechanism of IL-13-induced GSH depletion in airway epithelial cells, GSH depletion has been shown to promote T2 responses during chronic respiratory diseases. Kato et al. observed endocrine disrupting chemicals induced both GSH depletion and reciprocal regulation of IL-10/IL-12 production in airway epithelial cells.^74^ Reductive redox status, marked by high levels of GSH content, has been shown to promote IL-12 production via MAPK signaling in monocytes,^75^ whereas oxidative redox status, marked by low levels of GSH content, has been shown to promote IL-10 production via NF-κB activation in monocytes, B cells, T cells, and NK cells.^76,77^ No studies have confirmed these mechanisms of GSH depletion-induced T2 responses in airway epithelial cells. In the present study, we observed IL-13 induced both GSH depletion and T2 responses in the “T2-Skewed” phenotype. Yamanaka et al. observed IL-13 regulates IL-17C expression by inhibiting NF-κB activation in airway epithelial cells;^27^ and therefore, IL-13 may also induce GSH depletion and T2 responses by inhibiting NF-κB activation in the “T2-Skewed” phenotype. These hypotheses should be tested in future studies to better understand G×P effects on association, dissolution, impaired ion transport, and GSH depletion in relation to AgNP toxicity.

This is the first study to use organotypic cultures as a high-content *in vitro* model of the conducting airway to characterize G×P effects on AgNP toxicity, and therefore, refining this *in vitro* model in future studies is warranted. One limitation of this study was our use of only one sex and two genotypes; the use of both sexes and additional genotypes (e.g., BALB/cJ and SWR/J mice) or knockouts (e.g., *Nedd4l^-/-^* or *Nfkb1^-/-^ mice*) would increase genetic diversity and assist in defining a mechanistic basis for G×P effects on AgNP toxicity in future studies. A second limitation of this study was our use of IL-13 to skew differentiation toward an *in vitro* model of chronic respiratory diseases; the use of co-cultures with relevant cell populations, including mast cells or type 2 innate lymphoid cells (ILC2), would increase cellular diversity and improve the physiological relevance of this *in vitro* model in future studies. A third limitation of this study was our administration of the nominal dose of AgNP using suspensions in defined differentiation media. This approach may have contributed to their dissolution; the use of a Nano Aerosol Chamber for *in vitro* Toxicity studies (NACIVT) would allow for direct deposition of ENM onto the apical surface of organotypic cultures;^31,32,78^ or alternatively, the use of microfluidic systems would improve our understanding of the kinetics and dynamics of the conducing airway in future studies.^79^ Both approaches would allow this *in vitro* model to better recapitulate the physiological conditions of AgNP exposures in occupational settings.

Despite these limitations, this study establishes organotypic cultures derived from MTEC as a medium-throughput, high-content *in vitro* model for the conducing airway to characterize chemical perturbation as a means to screen and prioritize potential respiratory toxicants. Our results highlight the importance of considering dosimetry as well as G×P effects when screening and prioritizing potential respiratory toxicants. This is challenging and important for ENM, since their MoA have been shown to differ considerably but are still used in hundreds of consumer products. Prior to anticipating potential adverse organism responses arising from ENM toxicity, ensuring safe development of the consumer products in which they are used will be the most critical and necessary step toward safeguarding public health.

## List of Abbreviations

5×4 h: subacute exposure of 4 hours, every other day, over 5 days
Ag: silver mass
Ag^+^: silver ions
AgNP: silver nanoparticles
AJ or B6:Normal: A/J or C57BL6/J, “Normal” phenotype
AJ or B6:T2-Skewed: A/J or C57BL6/J, “T2-Skewed” phenotype
ALI: air-liquid interface
Au: gold mass
BMD: benchmark dose
BMDL: benchmark dose lower 95% confidence limit
Cd: cadmium mass
*Clca1*: calcium-activated chloride channel 1
COPD: chronic obstructive pulmonary disease
DCF: 2,7-dichlorofluorescein
DIV: day *in vitro*
ENaC: amiloride-sensitive epithelial Na^+^ channel
ENM: engineered nanomaterials
G×E: gene × environment interactions
G×P: genotype × phenotype interactions
GSH: glutathione
HEC: human equivalent concentration
ICPMS: inductively coupled plasma mass spectrometry
IHC: immunohistochemistry
ILC2: type 2 innate lymphoid cells
LDH: lactate dehydrogenase
MDA: malondialdehyde
MoA: mode of action
MTEC: murine tracheal epithelial cells
*Muc5ac*: mucin 5AC
NACIVT: Nano Aerosol Chamber for *in vitro* Toxicity studies
NDA: 2,3-naphthalenedicarboxaldehyde
*Nedd4l*: E3 ubiquitin-protein ligase NEDD4-like
NRC: National Research Council
OEL: occupational exposure limit
OR: odds ratio
PBPK: physiologically-based pharmacokinetic/toxicokinetic
QD: quantum dots
qRT-PCR: quantitative reverse transcription polymerase chain reaction
ROS: reactive oxygen species
T1/T2: type 1/type 2
TEER: transepithelial electrical resistance
TSCA: Toxic Substances Control Act
US EPA: U.S. Environmental Protection Agency

---

Equation 1. Nominal and dosimetric BMD (BMDL). *Y_i_* = adverse cellular response, *A* = intercept, *B* = slope, *X_i_* = dose, *R_i_* = random effect of preparation date for *i*^th^ sample, *r* = variance for random effects, *s* = residual variance for fixed effects, *σ* = population variability.

**Table S1.**
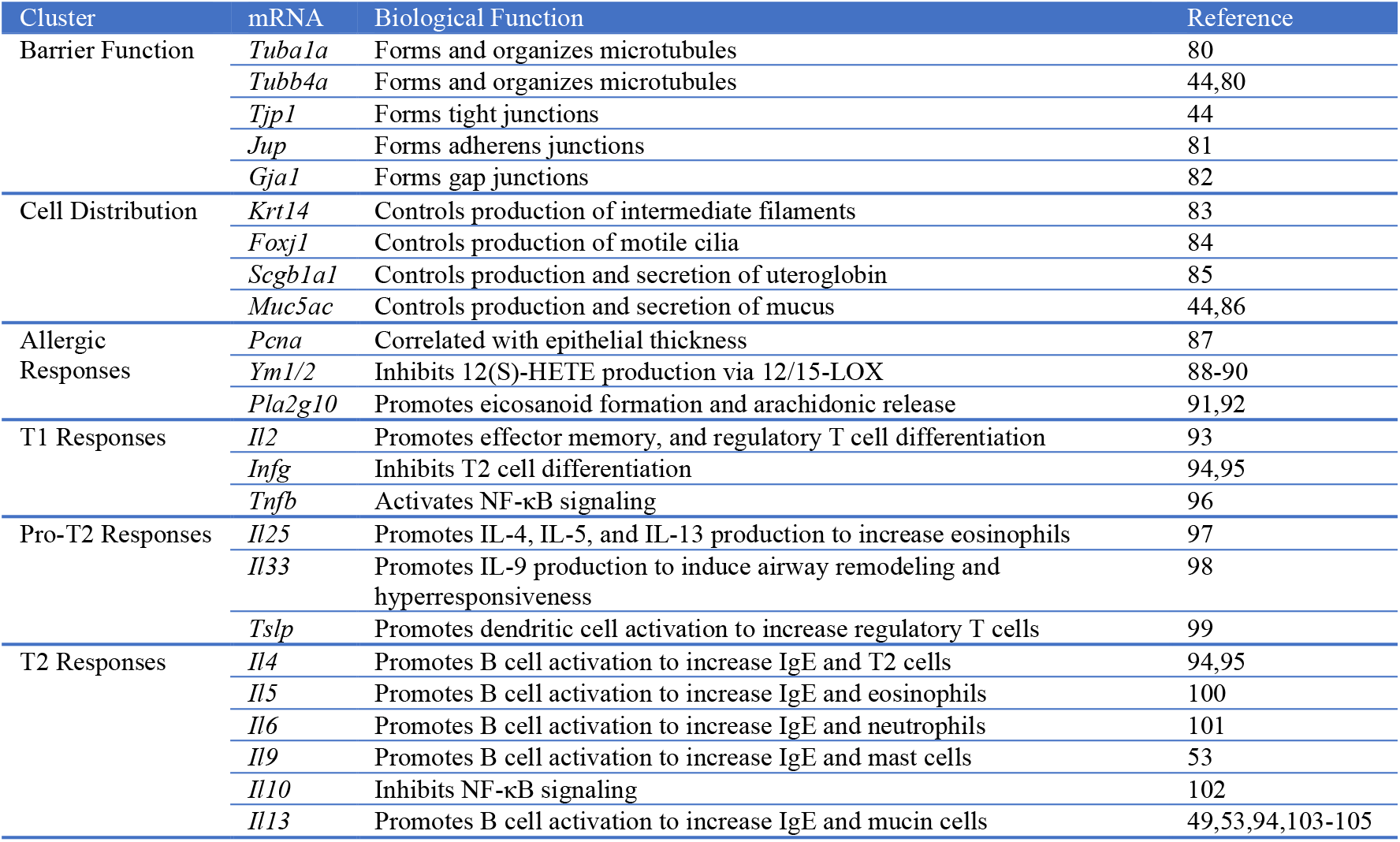
mRNA used for baseline characterization of organotypic cultures. mRNA: *Tuba1a* (tubulin alpha 1a); *Tubb4a* (tubulin beta 4A class IVa); *Tjp1* (tight junction protein 1); *Jup* (junction plakoglobin); *Gja1* (gap junction protein alpha 1); *Krt14* (keratin 14); *Foxj1* (forkhead box J1); *Scgb1a1* (secretoglobin family 1A member 1); *Muc5ac* (mucin 5AC); *Pcna* (proliferating cell nuclear antigen); *Ym1/2* (chitinase-3-like protein 1); *Pla2g10* (group 10 secretory phospholipase A2); *Il2* (interleukin 2); *Ifng* (interferon gamma); *Tnfb* (tumor necrosis factor beta); *Il25* (interleukin 25); *Il33* (interleukin 33); *Tslp* (thymic stromal lymphopoietin); *Il4* (interleukin 4); *Il5* (interleukin 5); *Il6* (interleukin 6); *Il9* (interleukin 9); *Il10* (interleukin 10); *Il13* (interleukin 13).

**Table S2.** Nominal dose of AgNP mass vs. dosimetric dose of Ag, Au, and Ag:Au mass. Exposure: h (hours). Genotype:Phenotype: AJ:Normal (A/J, “Normal” phenotype); B6:Normal (C57BL6/J, “Normal” phenotype); AJ:T2-Skewed (A/J, “T2-Skewed” phenotype); B6:T2-Skewed (C57BL6/J, “T2-Skewed” phenotype). Nominal Dose: AgNP (μg AgNP/mL media). Dosimetric Dose: Ag mass (μg Ag/mg protein); Au mass (μg Au/mg protein); Ag:Au mass (μg Ag:Au/mg protein). *n* = 3 biological replicates.

| Exposure | Genotype:Phenotype | Nominal Dose (AgNP) | Dosimetric Dose (Ag) | Dosimetric Dose Au) | Dosimetric Dose (Ag: Au) |
| --- | --- | --- | --- | --- | --- |
| 24 h | AJ:Normal | 0 | 0.00 | 0.00 | - |
|  |  | 12.5 | 4.03 | 0.75 | 5.41 |
|  |  | 25 | 9.43 | 1.06 | 8.90 |
|  |  | 50 | 21.07 | 4.00 | 5.27 |
|  | B6:Normal | 0 | 0.00 | 0.00 | - |
|  |  | 12.5 | 1.18 | 0.78 | 1.51 |
|  |  | 25 | 2.63 | 1.67 | 1.57 |
|  |  | 50 | 20.40 | 2.98 | 6.85 |
|  | AJ:T2-Skewed | 0 | 0.00 | 0.00 | - |
|  |  | 12.5 | 5.39 | 0.94 | 5.72 |
|  |  | 25 | 8.18 | 2.37 | 3.45 |
|  |  | 50 | 29.12 | 5.04 | 5.78 |
|  | B6:T2-Skewed | 0 | 0.04 | 0.00 | - |
|  |  | 12.5 | 7.34 | 0.31 | 23.76 |
|  |  | 25 | 15.94 | 0.70 | 22.81 |
|  |  | 50 | 29.25 | 5.91 | 4.95 |
| 5×4 h | AJ:Normal | 0 | 0.00 | 0.00 | - |
|  |  | 12.5 | 0.69 | 0.06 | 12.46 |
|  |  | 25 | 7.89 | 0.70 | 11.23 |
|  |  | 50 | 3.94 | 0.33 | 12.10 |
|  | B6:Normal | 0 | 0.01 | 0.00 | 22.80 |
|  |  | 12.5 | 1.20 | 0.10 | 11.74 |
|  |  | 25 | 1.93 | 0.17 | 11.46 |
|  |  | 50 | 3.79 | 0.33 | 11.36 |
|  | AJ:T2-Skewed | 0 | 0.00 | 0.00 | - |
|  |  | 12.5 | 0.37 | 0.09 | 4.08 |
|  |  | 25 | 0.26 | 0.06 | 4.12 |
|  |  | 50 | 7.86 | 1.87 | 4.21 |
|  | B6:T2-Skewed | 0 | 0.00 | 0.00 | - |
|  |  | 12.5 | 0.28 | 0.06 | 4.58 |
|  |  | 25 | 2.05 | 0.43 | 4.73 |
|  |  | 50 | 7.84 | 1.57 | 5.01 |

**Table S3.** Linear mixed effects model parameters of nominal dose of AgNP mass vs. dosimetric dose of Ag, Au, and Ag:Au mass. Dosimetric Dose: Ag mass (μg Ag/mg protein); Au mass (μg Au/mg protein); Ag:Au mass (μg Ag:Au/mg protein). Exposure: h (hours). Genotype:Phenotype: AJ:Normal (A/J, “Normal” phenotype); B6:Normal (C57BL6/J, “Normal” phenotype); AJ:T2-Skewed (A/J, “T2-Skewed” phenotype); B6:T2-Skewed (C57BL6/J, “T2-Skewed” phenotype). *n* = 3 biological replicates. Data represent mixed effects estimate of the linear slope parameters for each model; NS (*P*>0.05); * (*P*<0.05); ** (*P*<0.01); *** (*P*<0.001).

| Dosimetric Dose | Exposure | Genotype:Phenotype | Slope ($\beta$) | Intercept (B) | Correlation ($r^2$) | Significance ($P$) |
| --- | --- | --- | --- | --- | --- | --- |
| Ag | 24 h | AJ:Normal | 0.49 | -1.98 | 0.91 | *** |
|  |  | B6:Normal | 0.30 | -1.20 | 0.83 | *** |
|  |  | AJ:T2-Skewed | 0.68 | -2.76 | 0.92 | *** |
|  |  | B6:T2-Skewed | 0.66 | -0.12 | 0.98 | *** |
|  | 5×4 h | AJ:Normal | 0.06 | 0.31 | 0.24 | NS |
|  |  | B6:Normal | 0.08 | 0.10 | 0.99 | * |
|  |  | AJ:T2-Skewed | 0.19 | -1.44 | 0.82 | *** |
|  |  | B6:T2-Skewed | 0.18 | -1.22 | 0.87 | *** |
| Au | 24 h | AJ:Normal | 2.48E-04 | -9.33E-04 | 0.91 | *** |
|  |  | B6:Normal | 2.47E-04 | -6.67E-05 | 0.99 | *** |
|  |  | AJ:T2-Skewed | 4.91E-04 | -2.00E-03 | 0.91 | *** |
|  |  | B6:T2-Skewed | 3.49E-04 | -1.47E-03 | 0.81 | *** |
|  | 5×4 h | AJ:Normal | 1.52E-05 | 0.00E+00 | 0.29 | NS |
|  |  | B6:Normal | 2.29E-05 | -1.88E-19 | 0.73 | *** |
|  |  | AJ:T2-Skewed | 1.85E-04 | -1.47E-03 | 0.81 | *** |
|  |  | B6:T2-Skewed | 1.31E-04 | -8.67E-04 | 0.85 | *** |
| Ag:Au | 24 h | AJ:Normal | -0.01 | 6.40 | 0.00 | NS |
|  |  | B6:Normal | 0.02 | 3.23 | 0.02 | NS |
|  |  | AJ:T2-Skewed | 0.01 | 5.98 | 0.00 | NS |
|  |  | B6:T2-Skewed | -0.56 | 33.13 | 0.22 | NS |
|  | 5×4 h | AJ:Normal | 0.07 | 8.96 | 0.40 | * |
|  |  | B6:Normal | -0.19 | 18.46 | 0.44 | * |
|  |  | AJ:T2-Skewed | -0.03 | 5.14 | 0.30 | * |
|  |  | B6:T2-Skewed | 0.03 | 3.47 | 0.49 | ** |

**Table S4.**
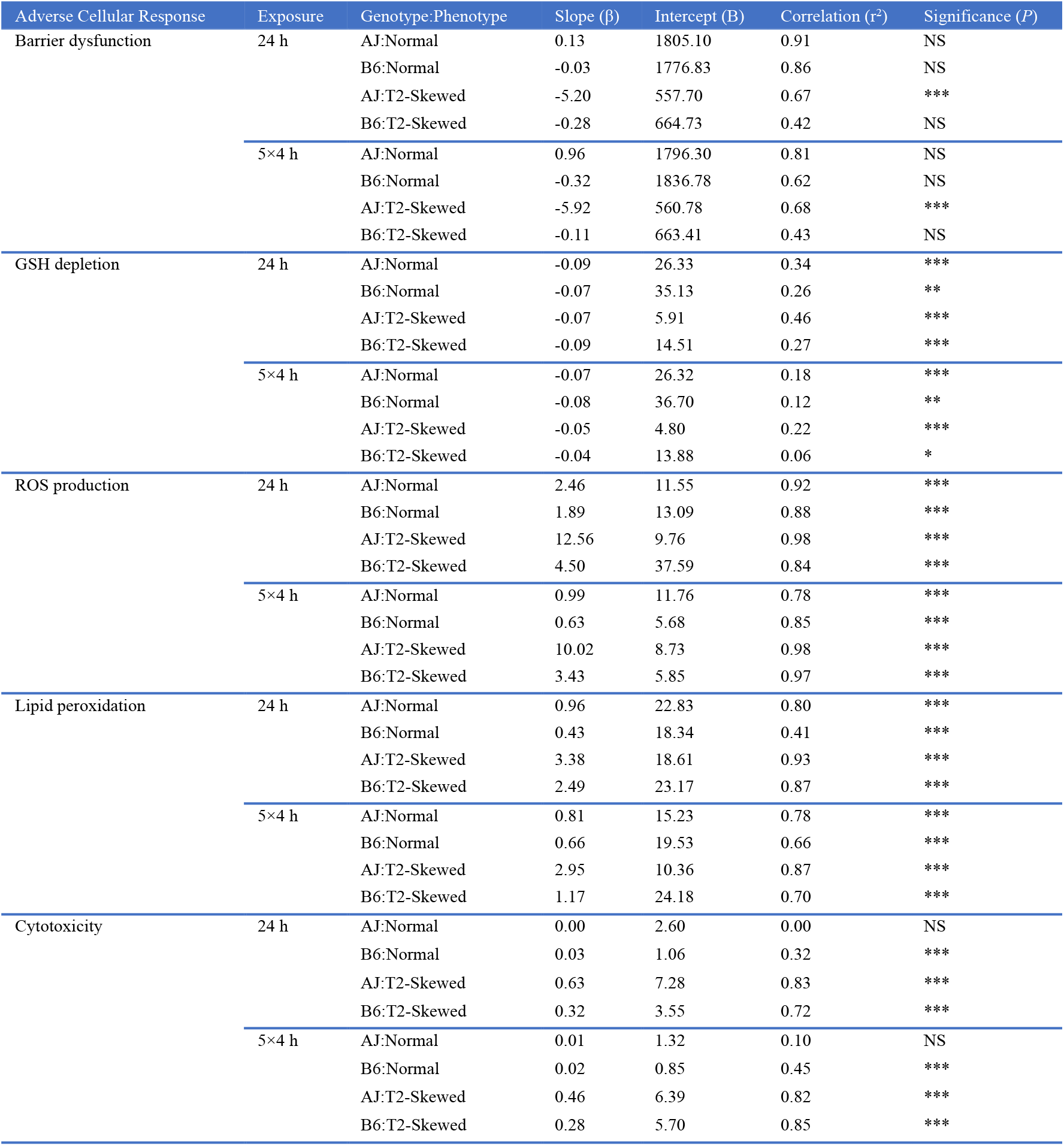
Linear mixed effects model parameters of nominal dose-response relationships. Exposure: h (hours). Genotype:Phenotype: AJ:Normal (A/J, “Normal” phenotype); B6:Normal (C57BL6/J, “Normal” phenotype); AJ:T2-Skewed (A/J, “T2-Skewed” phenotype); B6:T2-Skewed (C57BL6/J, “T2-Skewed” phenotype). *n* = 3 biological replicates with technical replicates. Data represent mixed effects estimate of the linear slope parameters for each model; NS (*P*>0.05); * (*P*<0.05); ** (*P*<0.01); *** (*P*<0.001).

**Table S5.**
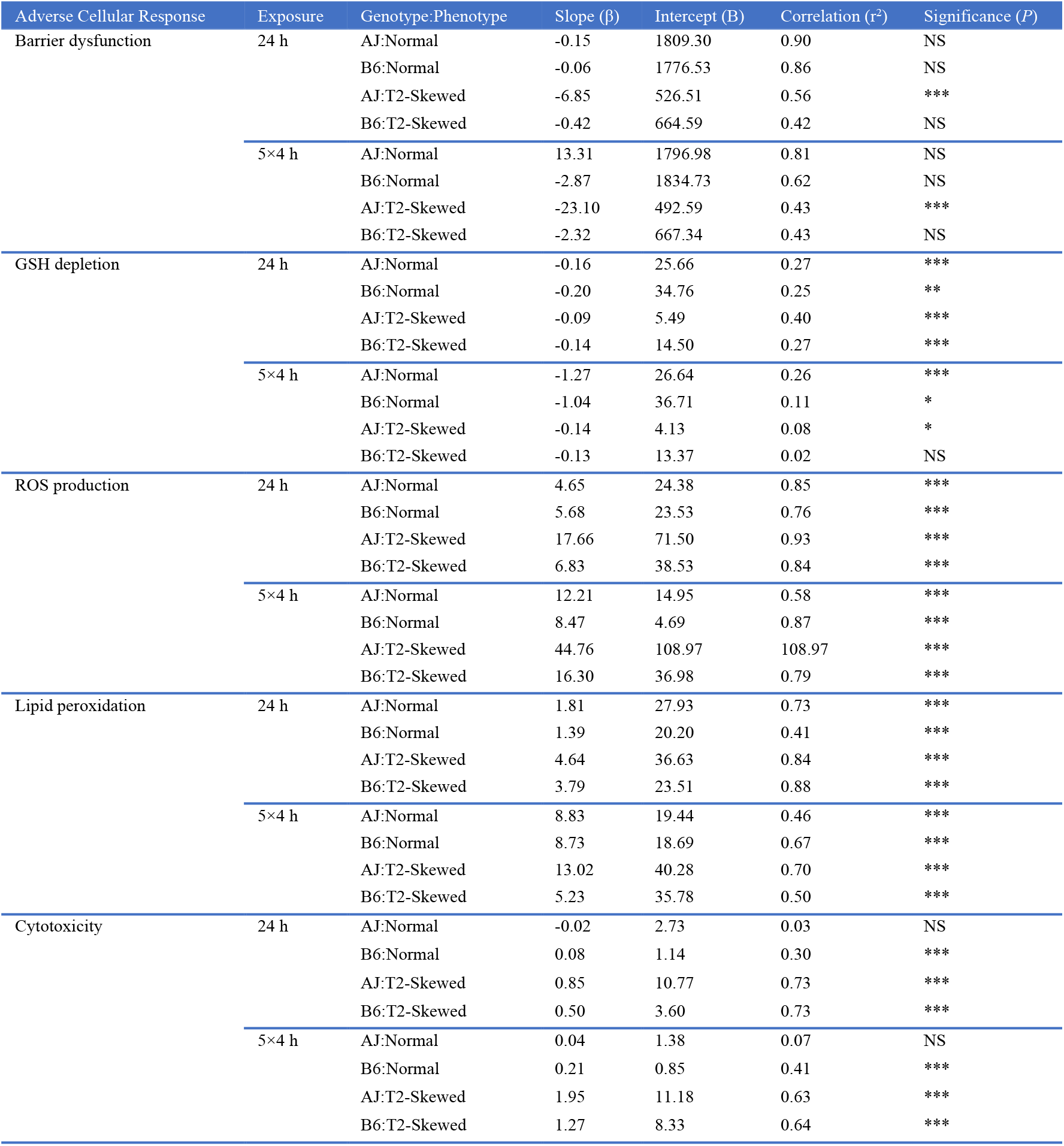
Linear mixed effects model parameters of dosimetric dose-response relationships. Exposure: h (hours). Genotype:Phenotype: AJ:Normal (A/J, “Normal” phenotype); B6:Normal (C57BL6/J, “Normal” phenotype); AJ:T2-Skewed (A/J, “T2-Skewed” phenotype); B6:T2-Skewed (C57BL6/J, “T2-Skewed” phenotype). *n* = 3 biological replicates with technical replicates. Data represent mixed effects estimate of the linear slope parameters for each model; NS (*P*>0.05); * (*P*<0.05); ** (*P*<0.01); *** (*P*<0.001).

**Table S6.** Nominal BMD (BMDL) vs. dosimetric BMD (BMDL) for adverse cellular responses. Exposure: h (hours). Genotype:Phenotype: AJ:Normal (A/J, “Normal” phenotype); B6:Normal (C57BL6/J, “Normal” phenotype); AJ:T2-Skewed (A/J, “T2-Skewed” phenotype); B6:T2-Skewed (C57BL6/J, “T2-Skewed” phenotype). Nominal BMD (BMDL): μg AgNP/mL media; Dosimetric BMD (BMDL): μg Ag/mg protein. *n* = 3 biological replicates with technical replicates. Data represent BMD (BMDL); NS (*P*>0.05); * (*P*<0.05); ** (*P*<0.01); *** (*P*<0.001).

| Adverse Cellular Response | Exposure | Genotype:Phenotype | Nominal BMD (BMDL) | Significance (*P*) | Dosimetric BMD (BMDL) | Significance (*P*) |
| --- | --- | --- | --- | --- | --- | --- |
| GSH depletion | 24 h | AJ:Normal | 27.40 (21.30) | *** | 17.40 (12.80) | *** |
|  |  | B6:Normal | 51.70 (33.50) | *** | 17.50 (11.00) | *** |
|  |  | AJ:T2-Skewed | 22.00 (17.90) | *** | 18.00 (14.10) | *** |
|  |  | B6:T2-Skewed | 35.50 (25.90) | *** | 20.00 (15.20) | *** |
|  | 5×4 h | AJ:Normal | 39.40 (27.90) | *** | 2.10 (1.60) | *** |
|  |  | B6:Normal | 55.60 (35.10) | *** | 4.50 (2.70) | *** |
|  |  | AJ:T2-Skewed | 35.50 (25.90) | *** | 12.80 (7.70) | *** |
|  |  | B6:T2-Skewed | 75.60 (42.20) | * | >8.70 | NS |
| ROS production | 24 h | AJ:Normal | 5.30 (5.00) | *** | 3.90 (3.60) | *** |
|  |  | B6:Normal | 6.70 (6.30) | *** | 3.20 (2.90) | *** |
|  |  | AJ:T2-Skewed | 2.60 (2.50) | *** | 3.50 (3.30) | *** |
|  |  | B6:T2-Skewed | 8.00 (7.40) | *** | 5.30 (4.90) | *** |
|  | 5×4 h | AJ:Normal | 9.90 (9.00) | *** | 1.10 (1.00) | *** |
|  |  | B6:Normal | 7.90 (7.30) | *** | 0.60 (0.50) | *** |
|  |  | AJ:T2-Skewed | 2.60 (2.50) | *** | 1.80 (1.60) | *** |
|  |  | B6:T2-Skewed | 3.30 (3.20) | *** | 1.80 (1.70) | *** |
| Lipid peroxidation | 24 h | AJ:Normal | 9.40 (8.50) | *** | 5.80 (5.20) | *** |
|  |  | B6:Normal | 21.00 (17.20) | *** | 6.50 (5.30) | *** |
|  |  | AJ:T2-Skewed | 5.20 (4.90) | *** | 5.60 (5.20) | *** |
|  |  | B6:T2-Skewed | 7.00 (6.50) | *** | 4.50 (4.20) | *** |
|  | 5×4 h | AJ:Normal | 9.90 (9.00) | *** | 1.40 (1.20) | *** |
|  |  | B6:Normal | 13.20 (11.60) | *** | 1.00 (0.90) | *** |
|  |  | AJ:T2-Skewed | 7.30 (6.80) | *** | 3.50 (3.00) | *** |
|  |  | B6:T2-Skewed | 11.90 (10.60) | *** | 8.30 (3.70) | *** |
| Cytotoxicity | 24 h | AJ:Normal | >50.00 | NS | >24.50 | NS |
|  |  | B6:Normal | 16.50 (14.10) | *** | 5.60 (4.70) | *** |
|  |  | AJ:T2-Skewed | 7.10 (6.60) | *** | >33.50 | NS |
|  |  | B6:T2-Skewed | 10.80 (9.70) | *** | 7.00 (6.30) | *** |
|  | 5×4 h | AJ:Normal | 55.40 (35.10) | *** | 8.30 (3.70) | *** |
|  |  | B6:Normal | 20.50 (16.90) | *** | 1.70 (1.40) | *** |
|  |  | AJ:T2-Skewed | 7.20 (6.70) | *** | 2.90 (2.50) | *** |
|  |  | B6:T2-Skewed | 7.80 (7.20) | *** | 2.60 (2.30) | *** |

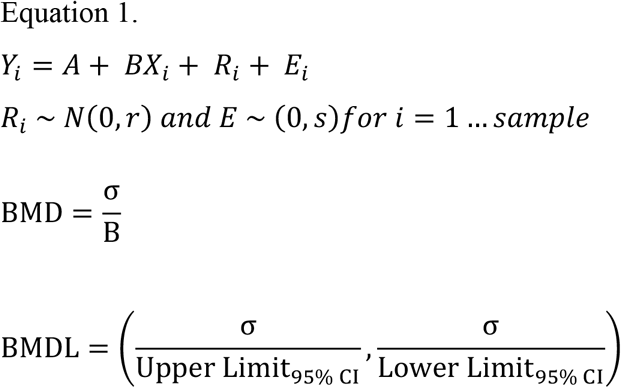

## References

1 Wilson, A. G. E. in New Horizons in Predictive Toxicology Drug Discovery 1–8 (2011).

2 Burden, N. et al. Aligning the 3Rs with new paradigms in the safety assessment of chemicals. Toxicology 330, 62–66, doi:10.1016/j.tox.2015.01.014 (2015).

3 Krewski, D., Andersen, M. E., Mantus, E. & Zeise, L. Toxicity testing in the 21st century: implications for human health risk assessment. Risk Anal 29, 474–479, doi:10.1111/j.1539-6924.2008.01150.x (2009).

4 Judson, R. S. et al. Estimating toxicity-related biological pathway altering doses for high-throughput chemical risk assessment. Chem Res Toxicol 24, 451–462, doi:10.1021/tx100428e (2011).

5 Hallstrand, T. S. et al. Airway epithelial regulation of pulmonary immune homeostasis and inflammation. Clinical Immunology 151, 1–15, doi:10.1016/j.clim.2013.12.003 (2014).

6 Weaver, M., Batts, L. & Hogan, B. Tissue interactions pattern the mesenchyme of the embryonic mouse lung. Developmental Biology 258, 169–184, doi:10.1016/s0012-1606(03)00117-9 (2003).

7 Rock, J. R. et al. Multiple stromal populations contribute to pulmonary fibrosis without evidence for epithelial to mesenchymal transition. Proceedings of the National Academy of Sciences 108, doi:10.1073/pnas.1117988108 (2011).

8 Borok, Z. et al. Cell Plasticity in Lung Injury and Repair. Proceedings of the American Thoracic Society 8, 215–222, doi:10.1513/pats.201012-067cb (2011).

9 Ostedgaard, L. S. et al. Gel-forming mucins form distinct morphologic structures in airways. Proc Natl Acad Sci U S A 114, 6842–6847, doi:10.1073/pnas.1703228114 (2017).

10 Steelant, B. et al. Histamine and T helper cytokine-driven epithelial barrier dysfunction in allergic rhinitis. J Allergy Clin Immunol 141, 951–963 e958, doi:10.1016/j.jaci.2017.08.039 (2018).

11 Flynn, A. N., Itani, O. A., Moninger, T. O. & Welsh, M. J. Acute regulation of tight junction ion selectivity in human airway epithelia. Proc Natl Acad Sci U S A 106, 3591–3596, doi:10.1073/pnas.0813393106 (2009).

12 MacRedmond, R. E. et al. Fluticasone Induces Epithelial Injury and Alters Barrier Function in Normal Subjects. J Steroids Horm Sci 5, doi:10.4172/2157-7536.1000134 (2014).

13 Matsui, H., Randell, S. H., Peretti, S. W., Davis, C. W. & Boucher, R. C. Coordinated clearance of periciliary liquid and mucus from airway surfaces. Journal of Clinical Investigation 102, 1125–1131, doi:10.1172/jci2687 (1998).

14 MacPherson, J. C. et al. Eosinophils Are a Major Source of Nitric Oxide Derived Oxidants in Severe Asthma: Characterization of Pathways Available to Eosinophils for Generating Reactive Nitrogen Species. The Journal of Immunology 166, 5763–5772, doi:10.4049/jimmunol.166.9.5763 (2001).

15 Donaldson, K., Stone, V., Clouter, A., Renwick, L. & MacNee, W. Ultrafine particles. Occup Environ Med 58, 211–216, 199 (2001).

16 Goss, C. H., Newsom, S. A., Schildcrout, J. S., Sheppard, L. & Kaufman, J. D. Effect of ambient air pollution on pulmonary exacerbations and lung function incystic fibrosis. Am J Respir Crit Care Med 169, 816–821, doi:10.1164/rccm.200306-779OC (2004).

17 Pope, C. A., 3rd. Epidemiology of fine particulate air pollution and human health: biologic mechanisms and who’s at risk? Environ Health Perspect 108 Suppl 4, 713–723 (2000).

18 USEPA. Reviewing New Chemicals under the Toxic Substances Control Act (TSCA), <https://www.epa.gov/reviewing-new-chemicals-under-toxic-substances-control-act-tsca/control-nanoscale-materials-under> (2017).

19 Judson, R. S. et al. In vitro screening of environmental chemicals for targeted testing prioritization: the ToxCast project. Environ Health Perspect 118, 485–492, doi:10.1289/ehp.0901392 (2010).

20 USEPA. Use of High-Throughput Assays and Computational Tools in the Endocrine Disruptor Screening Program, <https://www.epa.gov/endocrine-disruption/use-high-throughput-assays-and-computational-tools-endocrine-disruptor> (2013).

21 Quadros, M. E. & Marr, L. C. Silver Nanoparticles and Total Aerosols Emitted by Nanotechnology-Related Consumer Spray Products. Environmental Science & Technology 45, 10713–10719, doi:10.1021/es202770m (2011).

22 Quadros, M. E. & Marr, L. C. Environmental and Human Health Risks of Aerosolized Silver Nanoparticles. Journal of the Air & Waste Management Association 60, 770–781, doi:10.3155/1047-3289.60.7.770 (2010).

23 Vance, M. E. et al. Nanotechnology in the real world: Redeveloping the nanomaterial consumer products inventory. Beilstein journal of nanotechnology 6, 1769–1780, doi:10.3762/bjnano.6.181 (2015).

24 Kim, J. et al. Antimicrobial effects of silver nanoparticles. Nanomedicine: Nanotechnology, Biology and Medicine 3, 95–101, doi:10.1016/j.nano.2006.12.001 (2007).

25 Danilczuk, M., Lund, A., Sadlo, J., Yamada, H. & Michalik, J. Conduction electron spin resonance of small silver particles. Spectrochimica Acta Part A: Molecular and Biomolecular Spectroscopy 63, 189–191, doi:10.1016/j.saa.2005.05.002 (2006).

26 Ma, R. et al. Size-Controlled Dissolution of Organic-Coated Silver Nanoparticles. Environmental Science & Technology 46, 752–759, doi:10.1021/es201686j (2012).

27 Yamanaka, K. et al. IL-13 regulates IL-17C expression by suppressing NF-kappaB-mediated transcriptional activation in airway epithelial cells. Biochem Biophys Res Commun 495, 1534–1540, doi:10.1016/j.bbrc.2017.11.207 (2018).

28 Kim, H., Kim, M., Lee, S., Oh, S. & Chung, K. Genotoxic effects of silver nanoparticles stimulated by oxidative stress in human normal bronchial epithelial (BEAS-2B) cells. Mutation Research/Genetic Toxicology and Environmental Mutagenesis 726, 129–135, doi:10.1016/j.mrgentox.2011.08.008 (2011).

29 Cronholm, P. et al. Intracellular Uptake and Toxicity of Ag and CuO Nanoparticles: A Comparison Between Nanoparticles and their Corresponding Metal Ions. Small 9, 970–982, doi:10.1002/smll.201201069 (2013).

30 Gliga, A. R., Skoglund, S., Wallinder, I., Fadeel, B. & Karlsson, H. L. Size-dependent cytotoxicity of silver nanoparticles in human lung cells: the role of cellular uptake, agglomeration and Ag release. Particle and Fibre Toxicology 11, 1–17, doi:10.1186/1743-8977-11-11 (2014).

31 Jeannet, N., Fierz, M., Kalberer, M., Burtscher, H. & Geiser, M. Nano Aerosol Chamber for In-Vitro Toxicity (NACIVT) studies. Nanotoxicology 9, 34–42, doi:10.3109/17435390.2014.886739 (2014).

32 Jeannet, N. et al. Acute toxicity of silver and carbon nanoaerosols to normal and cystic fibrosis human bronchial epithelial cells. Nanotoxicology 10, 1–13, doi:10.3109/17435390.2015.1049233 (2015).

33 Kim, H. et al. Silver nanoparticles induce p53-mediated apoptosis in human bronchial epithelial (BEAS-2B) cells. The Journal of Toxicological Sciences 39, 401–412, doi:10.2131/jts.39.401 (2014).

34 Schlinkert, P. et al. The oxidative potential of differently charged silver and gold nanoparticles on three human lung epithelial cell types. Journal of Nanobiotechnology 13, 1, doi:10.1186/s12951-014-0062-4 (2015).

35 Zhang, H. et al. Mammalian Cells Exhibit a Range of Sensitivities to Silver Nanoparticles that are Partially Explicable by Variations in Antioxidant Defense and Metallothionein Expression. Small 11, 3797–3805, doi:10.1002/smll.201500251 (2015).

36 Choo, W. et al. Long-term exposures to low doses of silver nanoparticles enhanced in vitro malignant cell transformation in non-tumorigenic BEAS-2B cells. Toxicology in Vitro 37, 41–49, doi:10.1016/j.tiv.2016.09.003 (2016).

37 Lerner, C. A., Sundar, I. K. & Rahman, I. Mitochondrial redox system, dynamics, and dysfunction in lung inflammaging and COPD. Int J Biochem Cell Biol 81, 294–306, doi:10.1016/j.biocel.2016.07.026 (2016).

38 Alsaleh, N. B. & Brown, J. M. Immune responses to engineered nanomaterials: Current understanding and challenges. Current Opinion in Toxicology 10, 8–14, doi:10.1016/j.cotox.2017.11.011 (2018).

39 von Mutius, E. Gene-environment interactions in asthma. J Allergy Clin Immunol 123, 3–11; quiz 12-13, doi:10.1016/j.jaci.2008.10.046 (2009).

40 Molfino, N. A. & Coyle, A. J. Gene-environment interactions in chronic obstructive pulmonary disease. Int J Chron Obstruct Pulmon Dis 3, 491–497 (2008).

41 Lee, V. et al. Amphiphilic polymer-coated CdSe/ZnS quantum dots induce pro-inflammatory cytokine expression in mouse lung epithelial cells and macrophages. Nanotoxicology 9, 336–343, doi:10.3109/17435390.2014.930532 (2015).

42 Scoville, D. K. et al. Susceptibility to quantum dot induced lung inflammation differs widely among the Collaborative Cross founder mouse strains. Toxicology and Applied Pharmacology 289, 240–250, doi:10.1016/j.taap.2015.09.019 (2015).

43 Scoville, D. K. et al. Genetic determinants of susceptibility to silver nanoparticle-induced acute lung inflammation in mice. The FASEB Journal 31, 4600–4611, doi:10.1096/fj.201700187r (2017).

44 You, Y., Richer, E. J., Huang, T. & Brody, S. L. Growth and differentiation of mouse tracheal epithelial cells: selection of a proliferative population. Am J Physiol Lung Cell Mol Physiol 283, L1315–1321, doi:10.1152/ajplung.00169.2002 (2002).

45 Petecchia, L. et al. Cytokines induce tight junction disassembly in airway cells via an EGFR-dependent MAPK/ERK1/2-pathway. Lab Invest 92, 1140–1148, doi:10.1038/labinvest.2012.67 (2012).

46 Melgert, B. N. et al. Female mice are more susceptible to the development of allergic airway inflammation than male mice. Clinical & Experimental Allergy 35, 1496–1503, doi:10.1111/j.1365-2222.2005.02362.x (2005).

47 Elenkov, I. J. Glucocorticoids and the Th1/Th2 balance. Ann N Y Acad Sci 1024, 138–146, doi:10.1196/annals.1321.010 (2004).

48 Krause, S. et al. Blockade of interleukin-13-mediated cell activation by a novel inhibitory antibody to human IL-13 receptor alpha1. Mol Immunol 43, 1799–1807, doi:10.1016/j.molimm.2005.11.001 (2006).

49 Dickinson, J. D. et al. IL13 activates autophagy to regulate secretion in airway epithelial cells. Autophagy 12, 397–409, doi:10.1080/15548627.2015.1056967 (2016).

50 You, Y. & Brody, S. L. Culture and differentiation of mouse tracheal epithelial cells. Methods Mol Biol 945, 123–143, doi:10.1007/978-1-62703-125-7_9 (2013).

51 Horani, A., Dickinson, J. D. & Brody, S. L. Applications of mouse airway epithelial cell culture for asthma research. Methods Mol Biol 1032, 91–107, doi:10.1007/978-1-62703-496-8_7 (2013).

52 Park, J. J. et al. Characterization of 3D embryonic C57BL/6 and A/J mouse midbrain micromass in vitro culture systems for developmental neurotoxicity testing. Toxicol In Vitro 48, 33–44, doi:10.1016/j.tiv.2017.12.009 (2018).

53 Parker, J. C. et al. Chronic IL9 and IL-13 exposure leads to an altered differentiation of ciliated cells in a well-differentiated paediatric bronchial epithelial cell model. PLoS One 8, e61023, doi:10.1371/journal.pone.0061023 (2013).

54 Weldy, C. S. et al. Heterozygosity in the glutathione synthesis gene Gclm increases sensitivity to diesel exhaust particulate induced lung inflammation in mice. Inhal Toxicol 23, 724–735, doi:10.3109/08958378.2011.608095 (2011).

55 Juarez-Moreno, K. et al. Comparison of cytotoxicity and genotoxicity effects of silver nanoparticles on human cervix and breast cancer cell lines. Hum Exp Toxicol 36, 931–948, doi:10.1177/0960327116675206 (2017).

56 Long, Y. M. et al. Surface ligand controls silver ion release of nanosilver and its antibacterial activity against Escherichia coli. Int J Nanomedicine 12, 3193–3206, doi:10.2147/IJN.S132327 (2017).

57 Weldon, B. A. et al. Using primary organotypic mouse midbrain cultures to examine developmental neurotoxicity of silver nanoparticles across two genetic strains. Toxicol Appl Pharmacol, doi:10.1016/j.taap.2018.04.017 (2018).

58 Team, R. C. R: A language and environment for statistical computing, <https://www.r-project.org> (2018).

59 Gereda, J. E. et al. Relation between house-dust endotoxin exposure, type 1 T-cell development, and allergen sensitisation in infants at high risk of asthma. The Lancet 355, 1680–1683, doi:10.1016/s0140-6736(00)02239-x (2000).

60 Park, J. et al. Characterization of exposure to silver nanoparticles in a manufacturing facility. Journal of Nanoparticle Research 11, 1705–1712, doi:10.1007/s11051-009-9725-8 (2009).

61 Lee, J. et al. Exposure assessment of workplaces manufacturing nanosized TiO2 and silver. Inhalation Toxicology 23, 226–236, doi:10.3109/08958378.2011.562567 (2011).

62 Lee, J., Mun, J., Park, J. & Yu, I. A health surveillance case study on workers who manufacture silver nanomaterials. Nanotoxicology 6, 667–669, doi:10.3109/17435390.2011.600840 (2012).

63 Kim, E. et al. Case Study on Risk Evaluation of Silver Nanoparticle Exposure from Antibacterial Sprays Containing Silver Nanoparticles. Journal of Nanomaterials 2015, 1–8, doi:10.1155/2015/346586 (2015).

64 Weldon, B. A. et al. Occupational exposure limit for silver nanoparticles: considerations on the derivation of a general health-based value. Nanotoxicology 10, 1–13, doi:10.3109/17435390.2016.1148793 (2016).

65 Bachler, G., von Goetz, N. & Hungerbuhler, K. A physiologically based pharmacokinetic model for ionic silver and silver nanoparticles. Int J Nanomedicine 8, 3365–3382, doi:10.2147/IJN.S46624 (2013).

66 Soutiere, S. E., Tankersley, C. G. & Mitzner, W. Differences in alveolar size in inbred mouse strains. Respiratory Physiology & Neurobiology 140, 283–291, doi:10.1016/j.resp.2004.02.003 (2004).

67 Metzger, R. J., Klein, O. D., Martin, G. R. & Krasnow, M. A. The branching programme of mouse lung development. Nature 453, doi:10.1038/nature07005 (2008).

68 Chuang, H.-C. et al. Allergenicity and toxicology of inhaled silver nanoparticles in allergen-provocation mice models. International Journal of Nanomedicine 8, 4495–4506, doi:10.2147/ijn.s52239 (2013).

69 Alessandrini, F. et al. Pro-Inflammatory versus Immunomodulatory Effects of Silver Nanoparticles in the Lung: The Critical Role of Dose, Size and Surface Modification. Nanomaterials 7, 300, doi:10.3390/nano7100300 (2017).

70 Yasuo, M. et al. Relationship between calcium-activated chloride channel 1 and MUC5AC in goblet cell hyperplasia induced by interleukin-13 in human bronchial epithelial cells. Respiration 73, 347–359, doi:10.1159/000091391 (2006).

71 Kimura, T. et al. Deletion of the ubiquitin ligase Nedd4L in lung epithelia causes cystic fibrosis-like disease. Proc Natl Acad Sci U S A 108, 3216–3221, doi:10.1073/pnas.1010334108 (2011).

72 Campbell, C. D. et al. Whole-genome sequencing of individuals from a founder population identifies candidate genes for asthma. PLoS One 9, e104396, doi:10.1371/journal.pone.0104396 (2014).

73 Nam, J. H. et al. Interleukin-13/-4-induced oxidative stress contributes to death of hippocampal neurons in abeta1-42-treated hippocampus in vivo. Antioxid Redox Signal 16, 1369–1383, doi:10.1089/ars.2011.4175 (2012).

74 Kato, T. et al. Endocrine disruptors that deplete glutathione levels in APC promote Th2 polarization in mice leading to the exacerbation of airway inflammation. Eur J Immunol 36, 1199–1209, doi:10.1002/eji.200535140 (2006).

75 Utsugi, M. et al. Glutathione redox regulates lipopolysaccharide-induced IL-12 production through p38 mitogen-activated protein kinase activation in human monocytes: role of glutathione redox in IFN-gamma priming of IL-12 production. J Leukoc Biol 71, 339–347 (2002).

76 Haddad, J. J. & Fahlman, C. S. Redox-and oxidant-mediated regulation of interleukin-10: an anti-inflammatory, antioxidant cytokine? Biochem Biophys Res Commun 297, 163–176 (2002).

77 Meng, Q. et al. Nuclear Factor-kappaB modulates cellular glutathione and prevents oxidative stress in cancer cells. Cancer Lett 299, 45–53, doi:10.1016/j.canlet.2010.08.002 (2010).

78 Svensson, C. R. et al. Validation of an air-liquid interface toxicological set-up using Cu, Pd, and Ag well-characterized nanostructured aggregates and spheres. J Nanopart Res 18, 86, doi:10.1007/s11051-016-3389-y (2016).

79 Huh, D. et al. Reconstituting organ-level lung functions on a chip. Science 328, 1662–1668, doi:10.1126/science.1188302 (2010).

80 Marshall, C. B. et al. p73 Is Required for Multiciliogenesis and Regulates the Foxj1-Associated Gene Network. Cell Reports 14, 2289–2300, doi:10.1016/j.celrep.2016.02.035 (2016).

81 Smith, M. K., Koch, P. J. & Reynolds, S. D. Direct and indirect roles for beta-catenin in facultative basal progenitor cell differentiation. Am J Physiol Lung Cell Mol Physiol 302, L580–594, doi:10.1152/ajplung.00095.2011 (2012).

82 Martin, F. J. & Prince, A. S. TLR2 Regulates Gap Junction Intercellular Communication in Airway Cells. The Journal of Immunology 180, 4986–4993, doi:10.4049/jimmunol.180.7.4986 (2008).

83 Rock, J. R. et al. Basal cells as stem cells of the mouse trachea and human airway epithelium. Proc Natl Acad Sci U S A 106, 12771–12775, doi:10.1073/pnas.0906850106 (2009).

84 You, Y. et al. Role of f-box factor foxj1 in differentiation of ciliated airway epithelial cells. Am J Physiol Lung Cell Mol Physiol 286, L650–657, doi:10.1152/ajplung.00170.2003 (2004).

85 Wang, X. Y. et al. Novel method for isolation of murine clara cell secretory protein-expressing cells with traces of stemness. PLoS One 7, e43008, doi:10.1371/journal.pone.0043008 (2012).

86 Fahy, J. V. Goblet cell and mucin gene abnormalities in asthma. Chest 122, 320S–326S (2002).

87 Buchweitz, J. P., Harkema, J. R. & Kaminski, N. E. Time-dependent airway epithelial and inflammatory cell responses induced by influenza virus A/PR/8/34 in C57BL/6 mice. Toxicol Pathol 35, 424–435, doi:10.1080/01926230701302558 (2007).

88 Cai, Y., Kumar, R. K., Zhou, J., Foster, P. S. & Webb, D. C. Ym1/2 promotes Th2 cytokine expression by inhibiting 12/15(S)-lipoxygenase: identification of a novel pathway for regulating allergic inflammation. J Immunol 182, 5393–5399, doi:10.4049/jimmunol.0803874 (2009).

89 Ong, C. B. et al. Ozone-Induced Type 2 Immunity in Nasal Airways. Development and Lymphoid Cell Dependence in Mice. Am J Respir Cell Mol Biol 54, 331–340, doi:10.1165/rcmb.2015-0165OC (2016).

90 Chu, H. W. et al. Expression and activation of 15-lipoxygenase pathway in severe asthma: relationship to eosinophilic phenotype and collagen deposition. Clin Exp Allergy 32, 1558–1565 (2002).

91 Hallstrand, T. S. et al. Relationship between levels of secreted phospholipase A(2) groups IIA and X in the airways and asthma severity. Clin Exp Allergy 41, 801–810, doi:10.1111/j.1365-2222.2010.03676.x (2011).

92 Hallstrand, T. S. et al. Regulation and function of epithelial secreted phospholipase A2 group X in asthma. Am J Respir Crit Care Med 188, 42–50, doi:10.1164/rccm.201301-0084OC (2013).

93 Aoki, Y., Qiu, D., Uyei, A. & Kao, P. N. Human airway epithelial cells express interleukin-2 in vitro. Am J Physiol 272, L276–286, doi:10.1152/ajplung.1997.272.2.L276 (1997).

94 Ahdieh, M., Vandenbos, T. & Youakim, A. Lung epithelial barrier function and wound healing are decreased by IL-4 and IL-13 and enhanced by IFN-gamma. Am J Physiol Cell Physiol 281, C2029–2038, doi:10.1152/ajpcell.2001.281.6.C2029 (2001).

95 Saatian, B. et al. Interleukin-4 and interleukin-13 cause barrier dysfunction in human airway epithelial cells. Tissue Barriers 1, e24333, doi:10.4161/tisb.24333 (2013).

96 Neurath, M. F., Finotto, S. & Glimcher, L. H. The role of Th1/Th2 polarization in mucosal immunity. Nat Med 8, 567–573, doi:10.1038/nm0602-567 (2002).

97 Gordon, E. D., Locksley, R. M. & Fahy, J. V. Cross-Talk between Epithelial Cells and Type 2 Immune Signaling. The Role of IL-25. Am J Respir Crit Care Med 193, 935–936, doi:10.1164/rccm.201512-2534ED (2016).

98 Uchida, M. et al. Oxidative stress serves as a key checkpoint for IL-33 release by airway epithelium. Allergy 72, 1521–1531, doi:10.1111/all.13158 (2017).

99 Miyata, M. et al. Thymic stromal lymphopoietin is a critical mediator of IL-13-driven allergic inflammation. Eur J Immunol 39, 3078–3083, doi:10.1002/eji.200939302 (2009).

100 Salvi, S. et al. Interleukin-5 production by human airway epithelial cells. Am J Respir Cell Mol Biol 20, 984–991, doi:10.1165/ajrcmb.20.5.3463 (1999).

101 Xie, X. H. et al. Lipopolysaccharide induces IL-6 production in respiratory syncytial virus-infected airway epithelial cells through the toll-like receptor 4 signaling pathway. Pediatr Res 65, 156–162, doi:10.1203/PDR.0b013e318191f5c6 (2009).

102 Lim, S. et al. Differential expression of IL-10 receptor by epithelial cells and alveolar macrophages. Allergy 59, 505–514, doi:10.1111/j.1398-9995.2004.00455.x (2004).

103 Tyner, J. W. et al. Blocking airway mucous cell metaplasia by inhibiting EGFR antiapoptosis and IL-13 transdifferentiation signals. J Clin Invest 116, 309–321, doi:10.1172/JCI25167 (2006).

104 Thavagnanam, S. et al. Effects of IL-13 on mucociliary differentiation of pediatric asthmatic bronchial epithelial cells. Pediatr Res 69, 95–100, doi:10.1203/PDR.0b013e318204edb5 (2011).

105 Laoukili, J. et al. IL-13 alters mucociliary differentiation and ciliary beating of human respiratory epithelial cells. Journal of Clinical Investigation 108, 1817–1824, doi:10.1172/jci200113557 (2001).

